# Loss of genome maintenance accelerates podocyte damage and aging

**DOI:** 10.1101/2020.09.13.295303

**Authors:** Fabian Braun, Amrei M. Mandel, Linda Blomberg, Milagros N. Wong, Georgia Chatzinikolaou, Viji Nair, Roman Akbar-Haase, Victor G. Puelles, David H. Meyer, Phillip J. McCown, Fabian Haas, Mahdieh Rahmatollahi, Damian Fermin, Gisela G. Slaats, Tillmann Bork, Christoph Schell, Sybille Koehler, Paul T. Brinkoetter, Maja T. Lindenmeyer, Clemens D. Cohen, Martin Kann, Wilhelm Bloch, Matthew G. Sampson, Martijn ET Dollé, Matthias Kretzler, George A. Garinis, Tobias B. Huber, Bernhard Schermer, Thomas Benzing, Björn Schumacher, Christine E. Kurschat

## Abstract

DNA repair is essential for preserving genome integrity and ensuring cellular functionality and survival. Podocytes, post-mitotic glomerular epithelial cells, bear limited regenerative capacity, and their survival is indispensable to maintain the function of the kidney’s filtration units. While podocyte depletion is a hallmark of the aging process and of many proteinuric kidney diseases, the underlying factors remain unclear.

We investigated DNA repair in podocyte diseases by using a constitutive and an inducible podocyte-specific knockout mouse model for *Ercc1,* a multifunctional endonuclease cofactor involved in nucleotide excision repair (NER), interstrand crosslink (ICL) repair, and DNA double-strand break (DSB) repair. We assessed the consequences of *Ercc1* loss *in vivo,* complemented by mechanistical *in vitro* studies of induced DNA damage in cultured podocytes. Furthermore, we characterized DNA damage-related alterations in mouse and human renal tissue of different ages as well as in patient biopsies with minimal change disease and focal segmental glomerulosclerosis.

Podocyte-specific *Ercc1* knockout resulted in accumulation of DNA damage with ensuing proteinuria, podocyte loss, glomerulosclerosis, renal insufficiency, and reduced lifespan. The response to genomic stress was different to the pattern reported in other cell types, as podocytes activated mTORC1 signaling upon DNA damage *in vitro* and *in vivo*. The induced mTORC1 activation was abrogated by inhibiting DNA damage response through DNA-PK and ATM kinases *in vitro*. Moreover, pharmacological inhibition of mTORC1 modulated the development of glomerulosclerosis in *Ercc1*-deficient mice. Perturbed DNA repair gene expression and genomic stress was also detected in podocytes of human focal segmental glomerulosclerosis, characterized by podocyte loss. Beyond that, DNA damage accumulation occurred in podocytes of healthy aging mice and humans.

These findings reveal that genome maintenance is crucial for podocyte maintenance, linked to the mTORC1 pathway, and involved in the aging process as well as in the development of glomerulosclerosis, potentially serving as a therapeutic target in the future.

## Introduction

Most cells of the body are constantly subjected to various DNA damaging agents (1, 2). Therefore, cells depend on numerous DNA repair mechanisms to counteract transcription stress (3). Mutations in DNA repair genes result in a variety of pathologies ranging from cancer to progeroid syndromes (4, 5). The specific importance of genome maintenance in cells with limited regenerative capacity is demonstrated by the prevalence of neurodegeneration as a hallmark of DNA repair deficiency syndromes (6).

Glomerular podocytes are terminally differentiated, post-mitotic cells with little to no replacement capacity post-development (7). As an integral part of the primary filtration unit of the kidney (8), podocyte depletion is a leading cause of chronic kidney disease due to diabetes, hypertension, and other glomerulopathies, with ensuing loss of protein into the urine (9) and progressive renal insufficiency. A milder phenotype without overt proteinuria is present in aged kidneys, with globally sclerosed glomeruli due to decreasing numbers of podocytes with age (10). The precise pathomechanisms leading to podocyte depletion, however, are incompletely understood. Protecting this finite number of cells is, therefore, an important therapeutic goal (11).

Lately, first studies have indicated the importance of genome maintenance for renal health (12, 13). Mutations in the kinase endopeptidase and other proteins of small size (KEOPS) complex genes caused proteinuria and induced DNA damage response (DDR) *in vitro* (14). Likewise, glomerular DNA damage was found to be associated with declining kidney function (15) and cells isolated from the urine of patients suffering from diabetes and hypertension showed increased levels of DNA strand breaks (16). Furthermore, the glomeruli of a progeric mouse model were identified to share the expression profile of aged glomeruli, indicating a role of DNA damage repair in glomerular aging (17).

Several studies have proposed an interplay between DNA damage signaling and the mechanistic target of rapamycin (mTOR) pathway with factors induced by DNA damage exhibiting repressive effects on mTOR-complex1 (mTORC1) (18–20) and increased mTORC1 activity leading to transcription stress (21–23). This is of particular interest for podocytes, as they are highly dependent on a rigorous control of mTOR activity. While mTORC1-driven hypertrophy is a protective response upon podocyte depletion (10, 24, 25), mTORC1 overactivation drives pathologic hyperproliferation and sclerosis (26, 27). In line with these findings, side effects of pharmacological mTORC1 inhibition entail proteinuria and glomerular scarring (28–34).

Our group has proposed a role of DNA damage in glomerular aging by investigating glomeruli of the progeric *Ercc1* -/delta mouse (17). We followed up on these analyses and detected a newly discovered hallmark of aging – the shift towards transcription of smaller RNAs resulting from transcriptional stalling (35). To further address the involvement of podocytes in the process of accelerated aging, we created a podocyte-specific *Ercc1* knockout mouse model. Podocyte-specific DNA damage accumulation resulted in proteinuria, podocyte loss, glomerulosclerosis, and renal insufficiency in mice. Strikingly, both *in vivo* and *in vitro* analyses revealed mTORC1 activation upon DNA damage, indicating a cell type-specific response. Inhibiting DNA damage response through DNA-PK and ATM inhibition diminished mTORC1 activation. In turn, inhibiting mTORC1 activation moderately improved the glomerulosclerotic phenotypes in two Ercc1-deficient mouse models. In both humans and mice DNA damage foci were increased in podocyte nuclei of aged healthy kidneys accompanying mTORC1 activation. Ancillary, we identified perturbations in the expression of DNA repair genes in glomeruli and podocytes from human glomerular diseases associated with podocyte depletion and observed a significant increase of DNA damage in podocytes of patients suffering from focal segmental glomerulosclerosis. These results directly link genome maintenance to glomerular diseases and identify DNA damage accumulation as a hallmark of human podocyte aging.

## Methods

### GSEA Analysis

Raw data were downloaded from GSE43061, normalized with RMA (36), and further processed with limma (37). The log fold-changes of the comparison Ercc1 14 weeks vs. WT 14 weeks were used as input for the gene length GSEA analysis (17). The p-values and adjusted p-values are depicted in the plots. The enrichment analysis for chromosomal gene distributions was done in R v3.6.3 with the GSEA function of clusterProfiler (38), v3.14.3 was used with maxGSSize=6000 and nPerm=20000.

### Mice

Mice were bred in a mixed FVB/CD1 (*Ercc1* pko) or FVB/CD1/C57BL/6 (*Ercc1* ipko) background. All offspring was born in normal mendelian ratios. Mice were housed in the animal facility of the Center for Molecular Medicine Cologne or the Cluster of Excellence – Cellular Stress Responses in Aging-Associated Diseases. Following federal regulations, the Animal Care Committee of the University of Cologne reviewed and approved the experimental protocols. Animals were housed at specific pathogen-free (SPF) conditions with three-monthly monitoring according to FELASA suggestions. Housing was done in groups of less than six adult animals receiving CRM pelleted breeder and maintenance diet irradiated with 25 kGy (Special Diet Services, Witham, UK) and water *ad libitum*. Spot urine was collected once a week during cage changes or during sacrifice. Tamoxifen was administered at 400 mg/kg Tamoxifen in dry chow.

For rapamycin injection studies male and female *Ercc1*^fl/fl^ (ctrl) and *Ercc1*; *Nphs2.Cre* (pko) mice at week 6 of age were injected intraperitoneally 3 times/week with 2 mg/kg bodyweight of rapamycin diluted in 5% ethanol, 5% tween 80, and 5% PEG 400 or with 5% ethanol, 5% tween 80, and 5% PEG 400 as vehicle. Urine collection was performed 2 times/week and mice were sacrificed at week 13 of age for serum and kidney tissue isolation. All animals were maintained in grouped cages on a 12h light/dark cycle. Mice were kept on a regular diet and had access to water *ad libitum*. Body weight was measured weekly. Animals were housed in a temperature-controlled, pathogen-free animal facility at the Institute of Molecular Biology and Biotechnology (IMBB), which operates in compliance with the “Animal Welfare Act” of the Greek government, using the “Guide for the Care and Use of Laboratory Animals” as its standard.

Mice were anaesthetized by intraperitoneal injection of 10 µl per g bodyweight of 0.01% xylocaine and 12.5 mg/ml ketamine – blood was drawn from the left ventricle into a syringe rinsed with Heparin sulfate and animals were perfused with cold phosphate buffered saline (PBS). Kidneys were excised and embedded in OCT (Sakura, Torrance, CA) and frozen at -80°C or fixed in 4% neutral buffered formalin and subsequently embedded in paraffin.

Archival tissue of podocyte specific *Tsc*1 knockout mice was provided by Tillman Bork (University of Freiburg, Germany). *Tsc1*^fl/fl^ mice were crossbred with Nphs2-Cre^+^. For further details see ref (39).

Tissue of *Ercc1* -/delta mice was provided by Martin Dollé (National Institute of Public Health and the Environment Bilthoven, Netherlands). Further details provided in ref (40).

### Electron Microscopy

Mice were perfused with 4% paraformaldehyde and 2% glutaraldehyde in 0.1 M sodium cacodylate, pH 7.4. Postfixation was performed in the same buffer for two additional weeks at 4°C. Tissue was osmicated with 1% OsO4 in 0.1 M cacodylate and dehydrated in increasing ethanol concentrations. Upon infiltration and flat embedding were performed following standard procedures. Toluidine blue was used to stain semithin sections of 0.5 µm. 30 nm-thick sections were cut with an Ultracut UCT ultramicrotome (Reichert) and stained with 1% aqueous uranylic acetate and lead citrate. Samples were studied with Zeiss EM 902 and Zeiss EM 109 electron microscopes (Zeiss, Oberkochen, Germany).

### Podocyte isolation

To isolate primary podocytes, *Ercc1*^fl/fl^ mice heterozygous for the *R26mTmG* and *Nphs2.Cre* transgene were sacrificed and kidneys were used for glomerular preparation, as previously described (41). The glomeruli were digested and the single-cell suspension was used for fluorescence-activated cell sorting.

### qPCR Analysis

Total ribonucleic acid (RNA) was extracted from podocytes of *Ercc1/Nphs2.Cre/mTmG* mice using Direct-zol™ RNA MiniPrep Kit (cat. no. R2052, Zymo Research). Isolation of glomeruli, preparation of a glomerular single-cell suspension, and fluorescence-activated cell sorting was done as previously described (7). Podocytes were sorted into TriReagent (cat. no. 93289, Sigma-Aldrich). The complementary deoxyribonucleic acid (cDNA) was synthesized with High Capacity cDNA Reverse Transcription Kit (cat. no. 4368814, Applied Biosystems). PCR was performed using TaqMan™ Gene Expression Master Mix (cat. no. 4369016, Applied Biosystems) and the Applied Biosystems Real-time PCR system. Real-time PCR was measured with triplicates in each gene target. The sequence of the PCR primer used for *Ercc1* was: 5′-AGCCAGACCCTGAAAACAG-3′ and 5′-CACCTCACCGAATTCCCA-3′ in PrimeTime Mini qPCR Assay for *Ercc1* (Assay-ID: Mm.PT.58.42152282, IDT). The gene expression was calculated using comparative cycle threshold method and normalized to RNA polymerase II subunit A (*Polr2a*). The relative fold change of *Ercc1* expression in knockout mice was compared with WT and heterozygous mice.

### Urinary Albumin ELISA & Creatinine measurement

Urinary albumin levels were measured with a mouse albumin ELISA kit (mouse albumin ELISA kit; Bethyl Labs, Montgomery, TX, USA). Urinary creatinine kit (Cayman Chemical, Ann Arbor, MI, USA) was used to determine corresponding urinary creatinine values. For Coomassie Blue detection of albuminuria, spot urine of mice was diluted 1:20 in 1x Laemmli buffer and urinary proteins separated using poly-acrylamide gel electrophoresis with subsequent Coomassie gel stain.

### Plasma Creatinine and Urea measurement

Blood samples were centrifuged at 400 g 4°C for 20 minutes and plasma samples subsequently stored at -20°C until further analysis. Creatinine and Urea were measured using standard clinical protocols by the Department of Clinical Chemistry of the University of Cologne.

### Histologic analysis

To assess morphological changes in light microscopy we performed Periodic Acid Schiff staining. For specific antibody stainings, sections were deparaffinized in Xylene (VWR, Darmstadt, Germany), rehydrated in decreasing concentrations of ethanol, and subjected to heat-induced antigen retrieval in 10 mM Citrate Buffer pH 6 for 15 minutes. Peroxidase blocking was performed in methanol mixed with 3% hydrogen peroxidase (Roth, Karlsruhe, Germany) followed by Avidin/Biotin blocking (Vector, Burlingame, CA, USA) for 15 minutes each. After incubation in primary antibody (anti-phospho-S6 Ribosomal Protein (Ser235/236) # 4858 – Cell Signaling Technology) 1:200 in TBS 1% BSA at 4°C overnight, sections were washed in TBS and incubated in biotinylated secondary antibody (Jackson Immunoresearch, West Grove, USA) 1h at room temperature. For signal amplification the ABC Kit (Vector, Burlingame, CA, USA) was used before applying 3,30-diaminobenzamidine (Sigma-Aldrich, St Louis, USA) as a chromogen. Hematoxylin was used for counterstaining. After dehydration, slides were covered in Histomount (National Diagnostics, Atlanta, USA).

### Immunofluorescence Staining

Paraffin embedded tissue was cut into 3 µm thick sections and processed according to published protocols (42). Primary antibodies (anti-γH2A.X #2577s – Cell Signalling Technology, anti-nephrin #GP-N2 – Progen, anti-synaptopodin #65294 – Progen, anti-Dach1 #HPA012672 – Sigma Aldrich (43), anti-phospho-S6 Ribosomal Protein (Ser235/236) # 4858 – Cell Signaling Technology), and anti-p53 (anti-p53 #p53-protein-cm5 Leica Biosystems) were used at 1:200 dilution. Far-red fluorescent DNA dye Draq 5 was used as a nuclear marker.

Cells were processed according to published protocols (44).

For γH2A.X foci quantification, a custom-built FIJI macro was used. In brief, podocyte nuclei were identified through surrounding synaptopodin staining, segmented using the freehand tool and split into single channels. Draq 5 channel was converted into binary image using auto threshold “otsu dark” with subsequent particle measurement (range 5-infinite) to determine nuclear area. γH2A.X channel was converted into binary image using auto threshold “MaxEntropy dark” with subsequent particle measurement (range 0.02-infinite) to determine foci number and area.

### In vitro Experiments

Conditional immortalized murine podocytes were a gift by Stuart Shankland. Cells were cultured as previously described (45). Briefly, immortalized podocytes were cultured in RPMI media supplemented with 10% FBS and IFNγ (Sigma-Aldrich, Taufkirchen, Germany). Cells proliferated at 33°C on Primaria plastic plates (BD Biosciences, San Jose, CA, USA) until they reached a confluence of 60-70%. Differentiation of podocytes was induced by seeding the cells at 37°C in the absence of IFNy. After 10 days of differentiation, cells were treated with 5 or 10 µg/ml Mitomycin C (#M0503 - Sigma-Aldrich, Taufkirchen, Germany) for 2h in serum-free medium, followed by one washing step with PBS and another 6h incubation in serum-free medium without Mitomycin C before further processing. The absence of mycoplasm infection was tested regularly using the mycoplasm detection kit from Minerva biolabs (Minerva Biolabs, Berlin, Germany). For experiments with DNA damage response inhibitors, differentiated cells were pre-treated with 3 µM KU60019 (Selleckchem, Houston, TX, USA) or 1 µM Nedisertib (Selleckchem, Houston, TX, USA) for 1 h before inducing DNA damage by UV-C or Mitomycin C treatment. Inhibitors were added again after medium change following DNA damage induction to further incubate cells for 6 h before cell lysis.

### Western Blot analysis

SDS-PAGE was used for protein size separation with subsequent blotting onto polyvinylidene difluoride membranes and visualized with enhanced chemiluminescence after incubation of the blots with corresponding antibodies (Phospho-Histone H2A.X (Ser139); Phospho-S6 Ribosomal Protein (Ser235/236) (D57.2.2E); S6 Ribosomal Protein (5G10) – Cell Signaling Technology; alpha Actin – Developmental Studies Hybridoma Bank; beta-Tubulin (E7) - Developmental Studies Hybridoma Bank).

### ERCB Human microarray analysis

167 genes involved in DNA repair and nucleotide excision repair were compiled from the hallmark gene set “DNA-Repair” from the Molecular Signatures Database (MSigDB) Collection (46) and upon literature research. Human kidney biopsies and Affymetrix microarray expression data were obtained within the framework of the European Renal cDNA Bank - Kröner-Fresenius Biopsy Bank (47). Diagnostic biopsies were obtained from patients after informed consent and with approval of the local ethics committees. Following renal biopsy, the tissue was transferred to RNase inhibitor and micro-dissected into glomeruli and tubulo-interstitium. Total RNA was isolated, reverse transcribed, and amplified as previously reported (48). Fragmentation, hybridization, staining, and imaging were performed according to the Affymetrix Expression Analysis Technical Manual (Affymetrix, Santa Clara, CA, USA). Published datasets of glomerular samples were analysed for mRNA expression levels. Analysis included datasets from patients with minimal change disease (MCD; n=14), focal segmental glomerulosclerosis (FSGS; n=23), membranous nephropathy (MGN; n=21), IgA nephropathy (Glom; n=27), and hypertensive nephropathy (HTN; n=15) as well as controls (living donors (LD); n=42) (GSE99340, LD data from: GSE32591, GSE37463). CEL file normalization was performed with the Robust Multichip Average method using RMAExpress (Version 1.0.5) and the human Entrez-Gene custom CDF annotation from Brain Array version 18 (http://brainarray.mbni.med.umich.edu/Brainarray/default.asp). To identify differentially expressed genes, the SAM (Significance Analysis of Microarrays) method (49) was applied using SAM function in Multiple Experiment Viewer (TiGR MeV, Version 4.9). A q-value below 5% was considered to be statistically significant. The resulting gene expression list was censored for genes, whose products were detected in a transcriptomic and proteomic analysis of wild-type murine podocytes (41).

### Single nucleus sequencing

Nuclei were prepared from kidney biopsy cores stored in RNAlater from FSGS patients enrolled in the NEPTUNE study (50). The processing followed the protocol developed from the Kidney Precision Medicine Project. Nuclei preparations were processed and sequenced using 10x Genomics single cell sequencer. Analyses were performed on the output data files from CellRanger v6.0.0 using the Seurat R package (version 3.2 and 4.0; https://cran.r-project.org/web/packages/Seurat/index.html). To limit low quality nuclei and/or multiplets, we set gene counts and cutoffs to between 500 and 5000 genes and examined nuclei with a mitochondrial gene content of less than 10%. Nuclei were merged into a Seurat object using the CCA integrate function and nuclear cluster annotation was determined by finding enriched genes in each cell cluster. A comparison of these cluster selective gene profiles was compared against previously identified cell marker gene sets from human kidney samples from KPMP and other sources (50).

### Expression quantitative trait locus (eQTL) analysis

For the subgroup analysis of FSGS cohort, the procedure described in Gillies et al., 2018 was used with the following exceptions: only FSGS patients were analysed (N=87) and only RNAseq expression data for glomerular samples were utilized. Briefly, cis-eQTLs were identified using MatrixEQTL from among variants that were located either within the annotated boundaries of a gene or its surrounding region (+/-500 kb) (51). We then adjusted for age, sex, principal components of genetic ancestry, and the first 5 PEER factors (52). The genetic ancestry was calculated using LD-pruned WGS data from across all 87 patients using the EPACTS tool (https://genome.sph.umich.edu/wiki/EPACTS). The gene-level FDR for the MatrixEQTL was controlled using TORUS. Fine mapping of the eQTLs was performed using the DAP algorithm (53).

### Study approval

All investigations involving human specimen have been conducted according to the Declaration of Helsinki following approval of the local ethics committees. Written informed consent was received from participants prior to inclusion in the study. All mouse experiments were conducted according to institutional and federal guidelines and approved by the LANUV NRW VSG 84-02.04.2013.A336.

### Statistics

If not stated otherwise, unpaired two tailed Student’s t-test was used to compare two groups and p values ≤ 0.05 were considered significant. For multiple group comparisons, we applied 1-way ANOVA followed by Tukey’s post hoc correction. Statistics were performed using GraphPad Prism 8.

## Results

### DNA damage repair is essential for podocyte health

A well-established mouse model to induce DNA damage *in vivo* is the deletion of the DNA excision repair protein Ercc1, a cofactor for endonuclease Ercc4 (54–56). Our group has proposed a role of DNA damage in glomerular aging by investigating glomeruli of the *Ercc1* -/delta mouse (57), a model with whole body disruption of *Ercc1* on one allele and a truncated form of *Ercc1* on the second allele, leading to a hypomorphic variant with minimal residual Ercc1 activity. We previously showed that the expression profile of *Ercc1* -/delta glomeruli shared similarities with the profile of aged glomeruli. Very recently, a shift towards small RNA transcripts as a novel hallmark of aging was described (35), hence, we compared the length of RNA transcripts in prematurely aged 14-week-old *Ercc1* -/delta glomeruli in comparison to WT glomeruli of the same age. RNA expression showed an upregulation of the 5% and 1% shortest and downregulation of the 5% and 1% longest transcripts (Fig. 1A and S1) confirming transcriptional stalling as a hallmark of the aging process in *Ercc1*-deficient glomeruli (35). This glomerular phenotype coincided with the development of foot process effacement, a characteristic feature of podocyte damage, detected in electron microscopy (Fig. 1B), pointing towards DNA damage repair as an essential component of glomerular aging and podocyte health. To specifically investigate the role of DNA damage repair and aging on podocytes, we generated a constitutive podocyte-specific knockout of *Ercc1* using the cre-loxP system in mice of mixed FVB/CD1 background (58) (Fig. S2A & 1C.), expecting a phenotype of accelerated podocyte aging. Mice carrying the podocyte-specific *Ercc1* knockout (pko) had a dramatically decreased lifespan of 15-20 weeks, while cre negative animals (ctrl) and cre positive animals heterozygous for the floxed *Ercc1* allele (het) remained without overt abnormalities for up to 72 weeks (Fig. 1D & S2B & C).). This decreased lifespan was accompanied by significant albuminuria and severe renal failure represented by elevated serum creatinine and urea levels, starting at week 11 (Fig. 1E-G). At week 13, *Ercc1* pko mice developed severe generalized renal damage, including glomerulosclerosis, interstitial fibrosis, and tubular atrophy with protein casts (Fig. 1H).

**Figure 1:**
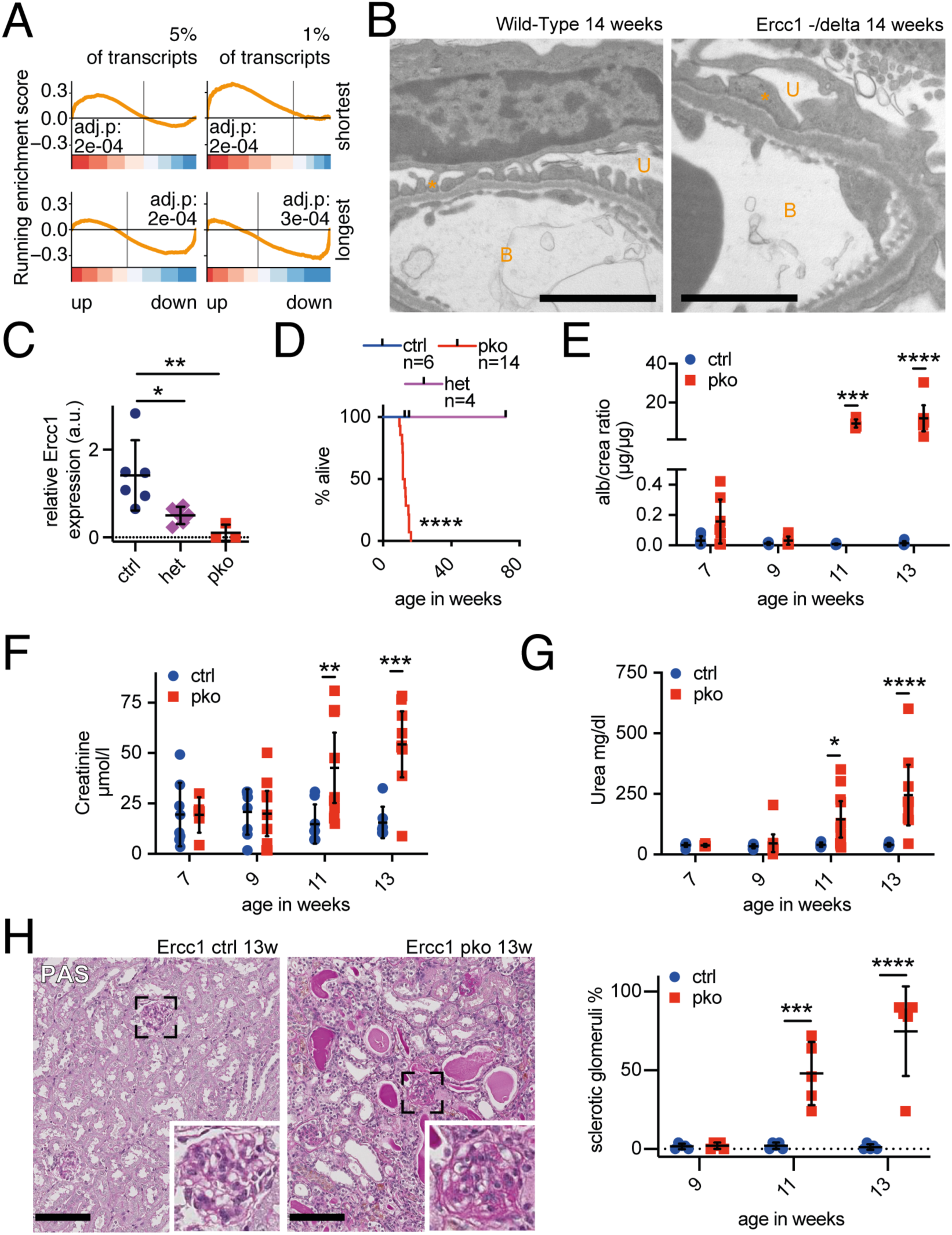
The podocyte-specific constitutive ko of *Ercc1* causes glomerulosclerosis. A: GSEA of gene classes according to transcript length. Shown are the analyses for the shortest 5 % (top left), shortest 1 % (top right), longest 5% (bottom left), and longest 1% (bottom right) of genes. The bottom color-coded panel shows the log2 fold changes of the microarray data in a ranked order. The top panels show the running enrichment score as an orange line. The smallest 1% (normalized enrichment score (NES) of 2.01) and 5% (NES of 1.37) of genes are significantly enriched in the upregulated genes, while the longest 1% (NES of -1.69) and 5% (NES of -1.42) of genes are significantly enriched in the downregulated genes in the comparison *Ercc1* -/delta 14 weeks vs. WT 14 weeks. B: Representative electron microscopy image of 14-week-old wild-type (WT) and *Ercc1* -/delta glomerular filtration barrier, scalebar indicating 2 µm, B: blood side – intracapillary space, U: urinary side – bowman’s space, asterisk indicating podocyte foot process (n=4). C: qPCR analysis for *Ercc1* in FACS-sorted podocytes of *Ercc1* ctrl, wt/pko (het) or pko mice. Delta-Delta-CT values expressed as scatterplots depicting mean plus 95% confidence interval. D: Kaplan-Meyer curve depicting survival of *Ercc1* ctrl, wt/pko (het) and pko mice (Mantel-Cox test). E: urinary albumin/creatinine analysis; F: serum creatinine analysis; G: serum urea analysis of *Ercc1* ctrl and pko mice. H: Representative Periodic Acid Schiff (PAS) staining of 13-week-old *Ercc1* ctrl and pko mice and quantification of sclerotic glomeruli, scalebars: 100 µm, n = 5, 50 glomeruli per sample. Scatterplots indicate mean plus 95% confidence interval, *p ≤ 0,05, **p ≤ 0,01, ***p ≤ 0,001, ****p ≤ 0,0001.

Similar results could be obtained in a tamoxifen-inducible podocyte-specific knockout (ipko) of *Ercc1* (Fig. S3), indicating that this phenotype was not dependent on developmental abnormalities in the absence of *Ercc1*.

### DNA damage accumulation in podocytes triggers cellular stress and podocyte loss

Ultrastructural alterations in 9-week-old constitutive *Ercc1* pko glomeruli were detectable in the form of focally effaced podocyte foot processes. These changes had not yet occurred in 7-week-old animals (Fig. 2A). Further evidence of podocyte stress was revealed by gradual reduction of nephrin abundance, an important protein of the glomerular filtration barrier, from weeks 9 to 13 (Fig. 2B & S4A). Podocyte number, glomerular hypertrophy, and podocyte density, investigated through staining of the podocyte specific proteins synaptopodin (SNP) and Dachshund Family Transcription Factor 1 (Dach1), remained within normal ranges at week 9 (Fig. 2C & S4B). In contrast, glomeruli from 11-week-old *Ercc1* pko mice clearly showed severe injury and loss of podocytes, indicated by the decrease and loss of both markers (Fig. 2D). This loss of podocytes was further validated through the analysis of WT1-positive cells in 9- and 11-week-old *Ercc1* pko mice (Fig. S4C).

**Figure 2:**
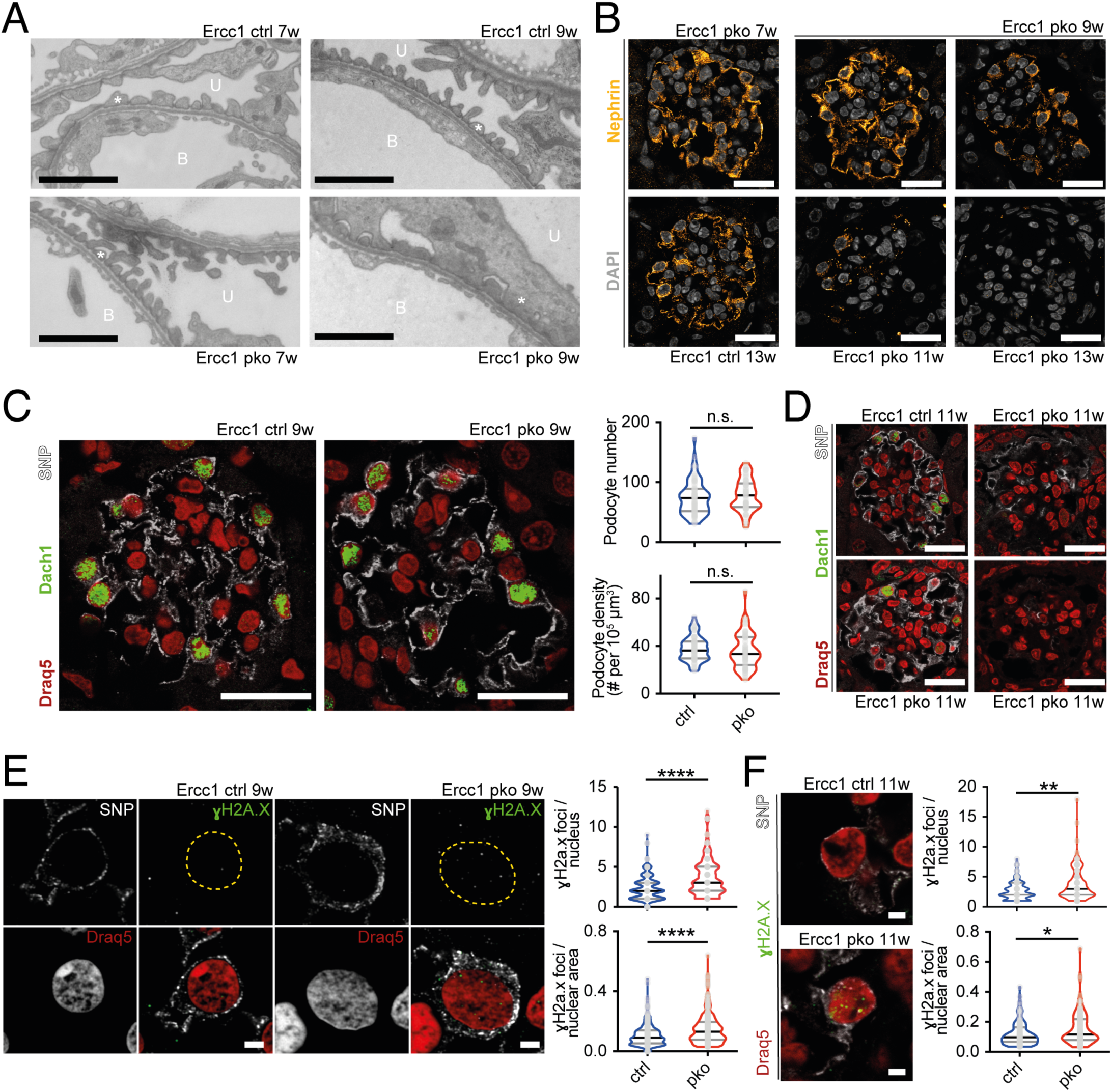
The podocyte-specific constitutive ko of *Ercc1* leads to foot process effacement and podocyte loss accompanied by accumulation of DNA damage. A: Representative electron microscopy image of 7- and 9-week-old *Ercc1* ctrl and pko slit diaphragms, scalebar indicating 2 µm, B: blood side – intracapillary space, U: urinary side – Bowman’s space, asterisk indicating podocyte foot process (n=3). B: Representative immunofluorescence staining of slit diaphragm protein nephrin (yellow) with nuclear marker DAPI (grey) of *Ercc1* ctrl at 13 weeks of age and pko kidneys at 7, 9, 11, and 13 weeks of age, scalebar indicating 2 µm (n=5). C: Representative immunofluorescence staining of podocyte proteins synaptopodin (SNP, gray), Dachshund homolog 1 (Dach1, green) (43) and far-red fluorescent DNA dye Draq5 (red) as a nuclear marker in sections of 9-week-old *Ercc1* ctrl and pko kidneys, with quantification of podocyte number and density of *Ercc1* ctrl and pko kidneys, scalebar indicating 10 µm (n=5). D: Corresponding staining of podocyte proteins synaptopodin (gray), Dachshund homolog 1 (Dach1, green) (43) and far-red fluorescent DNA dye Draq5 (red) as a nuclear marker in paraffin-embedded sections of 11-week-old *Ercc1* ctrl and pko kidneys, scalebar indicating 10 µm (n=5). E: Representative immunofluorescence staining of synaptopodin (SNP, gray), DNA damage marker γH2A.X (green) and nuclear marker Draq5 (red) in sections of 9-week-old *Ercc1* ctrl and pko kidneys, with quantification of γH2A.X foci per podocyte nucleus and nuclear area of *Ercc1* ctrl and pko kidneys, scalebar indicating 2 µm, yellow dotted line indicating nuclear border, n=5, 10 glomeruli per sample, 5 podocytes per glomerulus. F: Representative immunofluorescence staining of synaptopodin (SNP, gray), DNA damage marker γH2A.X (green) and nuclear marker Draq5 (red) in sections of 11-week-old *Ercc1* ctrl and pko kidneys, with quantification of γH2A.X foci per podocyte nucleus and nuclear area of *Ercc1* ctrl and pko kidneys, scalebar indicating 2 µm, n=5, 10 glomeruli per sample, 5 podocytes per glomerulus. All violin plots indicate median (black) and upper and lower quartile (gray), *p ≤ 0,05, **p ≤ 0,01, ***p ≤ 0,001, ****p ≤ 0,0001.

Foci of phosphorylated histone 2A.X (γH2A.X), a *bona fide* marker for DNA double-strand breaks, were significantly increased in both number and area in podocyte nuclei of *Ercc1* pko glomeruli at week 9 (Fig. 2E & S4D-F) compared to control animals. Strikingly, we also observed a smaller number of γH2A.X foci in almost all wildtype podocyte nuclei, indicative of constant DNA damage occurrence and subsequent repair in healthy glomeruli. At later time points, single podocytes with γH2A.X signals covering larger areas of the nuclei became apparent in *Ercc1* pko glomeruli (Fig. 2F).

### DNA damage accumulation in podocytes activates the mTORC1 pathway *in vivo*

Podocyte damage and loss is tightly linked to mTORC1 activation and cellular hypertrophy of the remaining podocytes (10, 24, 59). Therefore, we investigated the timepoint of mTORC1 activation in *Ercc1* pko mice. Interestingly, we detected a significant increase in pS6RP-positive cells in *Ercc1* pko glomeruli at 9 weeks of age (Fig. 3A), a timepoint when no evidence for podocyte loss was present yet (Fig. 2C & S4C). In-detail analysis revealed that more than 40% of podocytes showed mTORC1 activation at week 9 (Fig. 3A).

**Figure 3:**
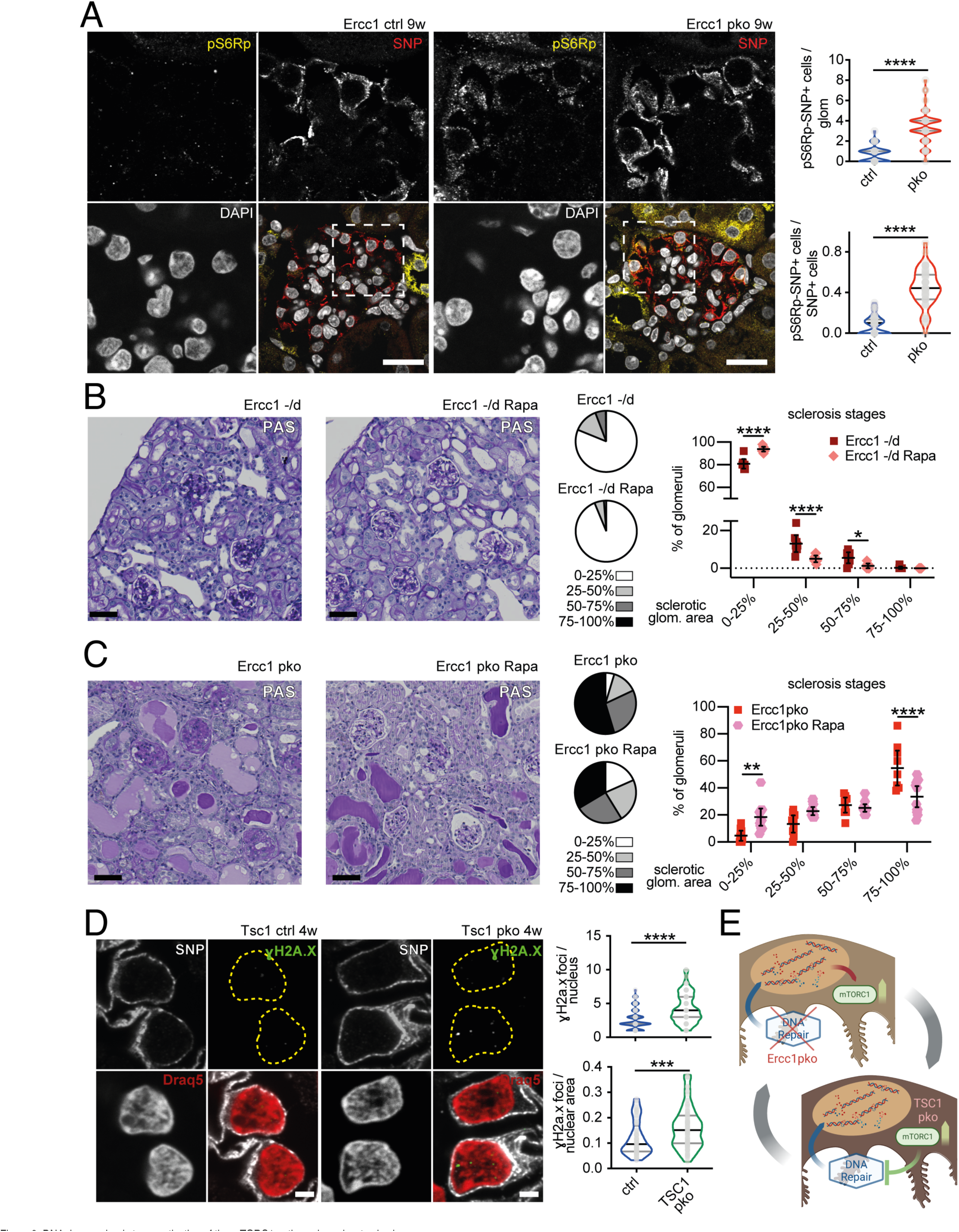
DNA damage leads to an activation of the mTORC1 pathway in podocytes *in vivo*. A: Representative immunofluorescence staining of SNP, pS6RP and DAPI in sections of 9-week-old *Ercc1* ctrl and pko kidneys with quantification of SNP and pS6RP double positive cells per glomerulus and per total SNP positive cells, scalebar indicating 10 µm (n=5, 10 glomeruli per sample). B: Representative Periodic Acid Schiff (PAS) staining of end-of-life *Ercc1* -/delta mice treated with 14 mg rapamycin per kg food from 8 weeks of age and glomerulosclerosis-assessment of all glomeruli depicted as parts of a whole and scatter plot (n = 8, 50 glomeruli per sample), scalebar indicating 50 µm. C: Representative Periodic Acid Schiff (PAS) staining of *Ercc1* pko mice treated with vehicle (*Ercc1* pko) or 2mg rapamycin per kg bodyweight (*Ercc1* pko Rapa) from 6 weeks of age and glomerulosclerosis-assessment of all glomeruli depicted as parts of a whole and scatter plot (n ≥ 9, 50 glomeruli per sample), scalebar indicating 50 µm. D: Representative immunofluorescence staining of synaptopodin (SNP, gray), DNA damage marker γH2A.X (green) and nuclear marker Draq5 (red) in sections of 4-week-old *Tsc1* ctrl and pko kidneys, with quantification of γH2A.X foci per podocyte nucleus and nuclear area of *Ercc1* ctrl and pko kidneys, yellow dotted line indicating nuclear border (n=5, 10 glomeruli per sample, 5 podocytes per glomerulus), scalebar indicating 2 µm. E: Schematic overview depicting the potential interplay between defective DNA damage repair and increased mTORC1 signaling. In *Ercc1* pko mice, accumulation of DNA damage triggers mTORC1 signaling. In *Tsc1* pko mice, hyperactive mTORC1 signaling also leads to increased DNA damage foci. All violin plots indicate median (black) and upper and lower quartile (gray), scatterplots indicate mean plus 95% confidence interval, *p ≤ 0,05, **p ≤ 0,01, ***p ≤ 0,001, ****p ≤ 0,0001.

To investigate whether increased mTORC1 signaling contributes to the development of podocyte loss, we analysed archival kidney tissue of *Ercc1* -/delta mice treated with 14 mg/kg food of the mTORC1 inhibitor rapamycin from 8 weeks of age until termination of the experiment due to high moribund scoring (40). Despite the fact that the treated cohort did not have an extended lifespan, moribund animals of the end-of-life cohort (aged 15-26 weeks) presented with a significant reduction of sclerotic glomeruli when treated with rapamycin (Fig. 3B). Functional data through blood and urine was not available for this cohort. In addition, we treated our podocyte-specific *Ercc1* ko mouse model with 2 mg/kg bodyweight rapamycin via intraperitoneal injections three times a week, beginning at 6 weeks of age. Again, we detected a modulation of the glomerular phenotype with a significant reduction in globally sclerotic glomeruli and an increase in mildly affected glomeruli upon rapamycin treatment at the end of our observation period at week 13 (Fig. 3C). Under these rapamycin conditions, albuminuria, serum creatinine, and serum urea levels were not reduced (Fig.S5A-C). However, rapamycin therapy was accompanied by a reduction of body weight to normal levels in treated animals (Fig.S5D & E), representative of a decreased rate of edema (Fig.S5F).

The observed detection of increased mTORC1 signaling and DNA damage accumulation suggests a potential interplay. It has been shown that increased mTORC1 signaling decreases the cellular ability to repair DNA damage (21, 22). Thus, we investigated the occurrence of increased DNA damage in a podocyte-specific *Tsc1* knockout mouse model, known to exhibit mTORC1 hyperactivation, by assessing γH2A.X foci accumulation in archival kidney tissue (10, 39). Indeed, *Tsc1* pko mice also depicted an increased number of DNA damage foci already at 4 weeks of age, when the phenotype is predominantly driven by mTORC1 hyperactivation (Fig. 3D). These data indicate that increased mTORC1 signaling due to DNA damage accumulation and decreased DNA damage repair upon mTORC1 activation may constitute a downward spiral aggravating podocyte injury (Fig. 3E).

### DNA damage activates the mTORC1 cascade through DNA damage signaling kinase DNA-PK in podocytes

To further investigate the mechanism behind mTORC1 activation occurring in podocytes upon transcription stress, we induced DNA damage through mitomycin C (MMC) treatment or UV-C irradiation *in vitro* in immortalized mouse podocytes (Fig. 4A). These treatments led to significant increases of γH2A.X (Fig. 4B) and accumulation of the DNA damage response protein p53 in the nucleus (Fig. 4C). Again, DNA damage induced an increased phosphorylation of S6 ribosomal protein (pS6RP), which is a downstream target of mTORC1, in podocytes (Fig. 4D). This phosphorylation was completely abrogated by mTORC1 inhibitor rapamycin and reduced by serum starvation, a well-known mTORC1 modulator (60). In order to identify the link between DNA damage accumulation and mTORC1 activation, we treated MMC-or UV-C-stimulated murine podocytes with inhibitors of the DNA damage signaling cascade (Fig. 4E). In both conditions, inhibition of DNA-dependent protein kinase (DNA-PK) by Nedisertib resulted in abrogation of S6RP phosphorylation. Similar results could be achieved with the Ataxia Telangiectasia Mutated serine/threonine kinase (ATM) inhibitor KU60019. No effects were detected upon treatment with the ATR inhibitor VE822 or with the CHK1 inhibitor prexasertib (data not shown). These data indicate a direct mechanistic link between transcription stress and mTORC1 signaling in podocytes.

**Figure 4:**
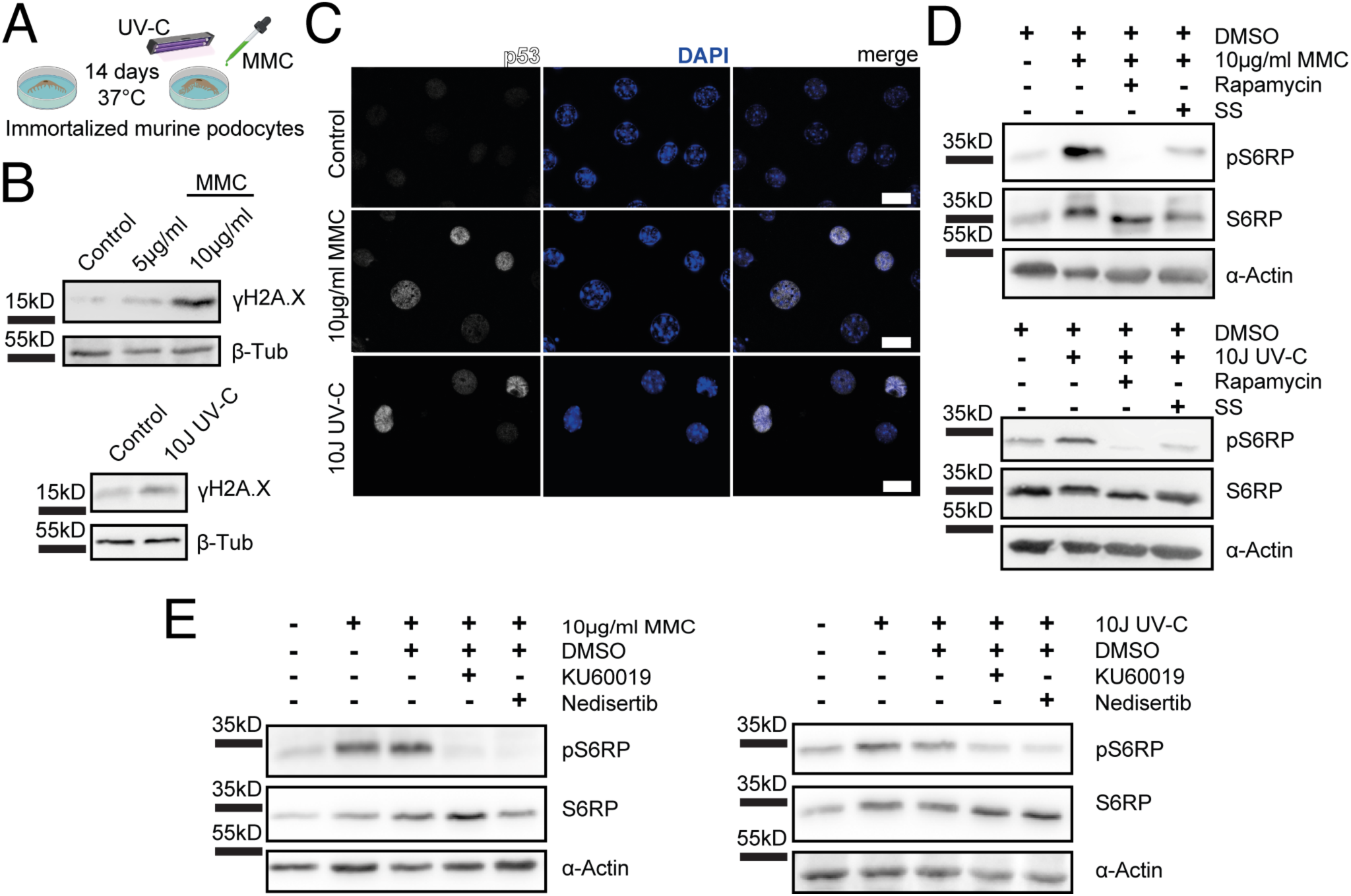
DNA damage leads to an activation of the mTORC1 pathway in podocytes through a DNA-PK-dependent mechanism. A: Schematic *in vitro* protocol for the induction of DNA damage. B: Representative immunoblot images for DNA damage marker γH2A.X and loading control protein beta-tubulin of immortalized murine podocyte lysates (n=3). C: Representative immunofluorescence images for tumor suppressor p53 and nuclear marker DAPI in immortalized murine podocytes (n=3). D: Representative immunoblot images for mTORC1 target phospho-S6 ribosomal protein (pS6RP), S6RP and loading control protein alpha-actin of immortalized murine podocyte lysates (n=3). All cells imaged or lysed after treatment with mitomycin C (MMC) or ultraviolet C (UV-C) irradiation ± 10 ng/ml rapamycin or serum starvation (SS), n ≥ 4. E: Representative immunoblot images for mTORC1 target phospho-S6 ribosomal protein (pS6RP), S6RP and loading control protein alpha-actin of immortalized murine podocyte lysates (n=3 MMC; n=6 UV-C). All cells imaged or lysed after treatment with mitomycin C (MMC) or ultraviolet C (UV-C) irradiation ± ATM inhibitor KU60019 or DNA-PK inhibitor nedisertib.

### DNA damage repair pathways are impaired in focal segmental glomerulosclerosis resulting in DNA damage accumulation

Since our initial goal was the analysis of podocyte aging, we investigated both murine and human glomeruli of young and aged subjects. Indeed, the well-documented increase in mTORC1 signaling in aged murine wild-type podocytes coincided with increased detection of DNA damage foci (Fig. 5A & S6A). Similar results were detected in a series of human healthy kidney tissue when comparing a young (21-49 years; n=4) with an aged group (69-81 years; n=5) (Fig.5B & S6B). These results confirm the occurrence of DNA damage in both healthy murine and human podocytes and indicate an association between DNA damage and podocyte aging, presumably contributing to age-related podocyte loss (10).

**Figure 5:**
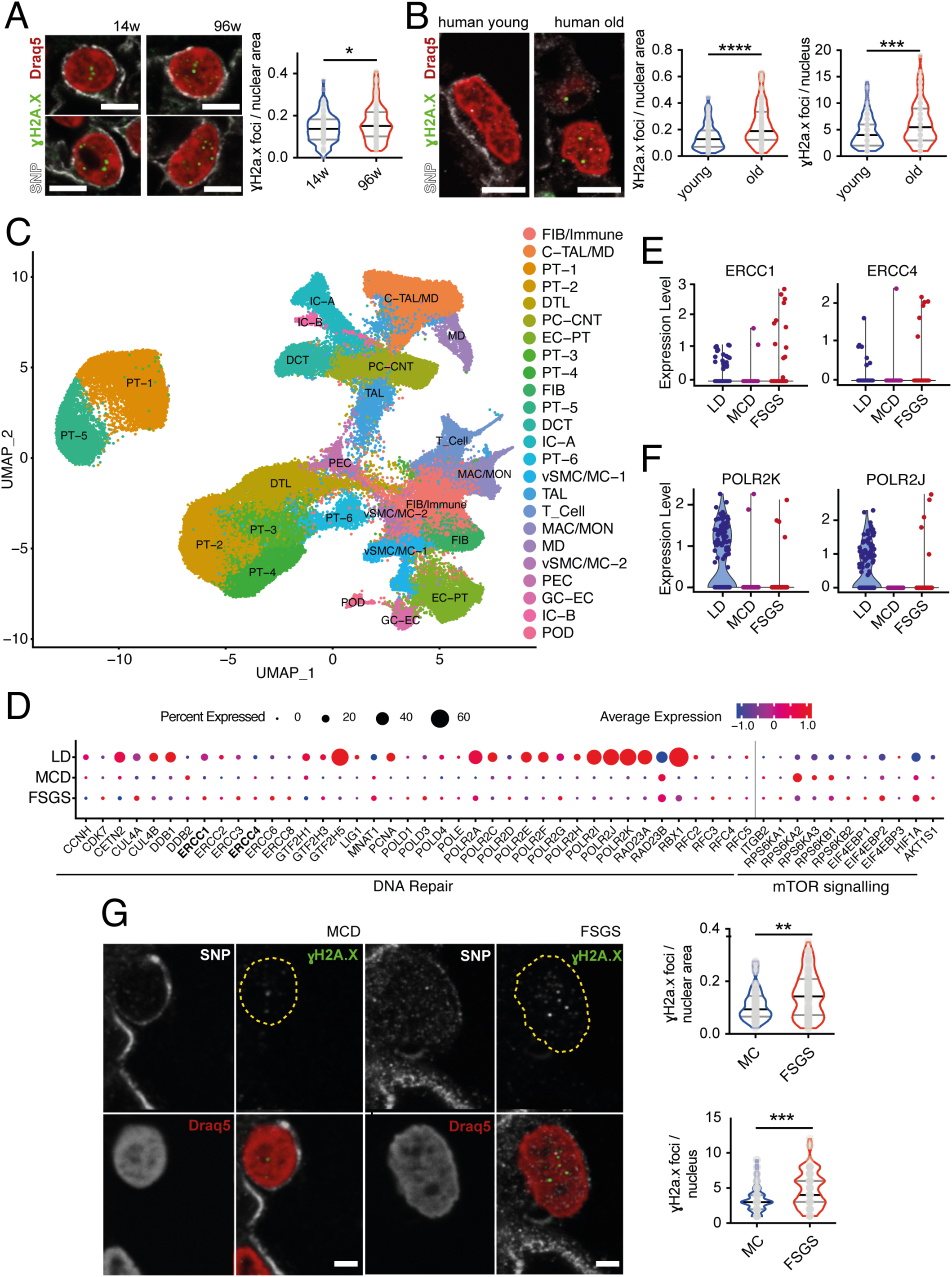
Podocytes accumulate DNA damage with aging and during human FSGS. A: Representative immunofluorescence staining of SNP, γH2A.X, and Draq5 in sections of murine young and aged wildtype kidneys with quantification of γH2A.X foci per podocyte nuclear area, scalebar indicating 2 µm, n=4, 5 glomeruli per sample, 5 podocytes per glomerulus. B: Representative immunofluorescence staining of SNP, γH2A.X, and Draq5 in sections of human young and old tumor nephrectomy kidneys with quantification of γH2A.X foci per podocyte nuclear area and per podocyte nucleus, scalebar indicating 5 µm, n≥4, 5 glomeruli per sample, 5 podocytes per glomerulus. C: UMAP of single nucleus sequencing data set (50) depicting the identification of 24 different cell types in analysed control, minimal change disease, and focal segmental glomerulosclerosis biopsies. FIB/Immue: Fibroblasts/Immune cells, C-TAL/MD: Thick ascending limb of loop of Henle, PT-1-6: proximal tubular cells, DTL: descending thin limb, PC-CNT: Connecting Tubule, EC-PT: Endothelial Cell, FIB: Fibroblast, DCT: Distal convoluted tubule, IC-A: Intercalated cell A, vSMC/MC-1: vascular smooth muscle cell/Muscle cell1, TAL: Thick ascending limb of loop of Henle, T_Cell: T cells, MAC/MON: Macrophage/Monocyte, MD: Medullary cell, vSMC/MC-2: vascular smooth muscle cell/Muscle cell2, PEC: Parietal epithelial cell, GC-EC: Glomerular endothelial cell, IC-B: Intercalated cell B, POD: Podocyte.. D: Bubble plot indicating the differences in DNA repair and mTORC1 target gene expression in podocytes between living donor (LD) kidney samples, Minimal change disease (MCD) and FSGS biopsies obtained through single nucleus sequencing (50). Bubble size indicating percentage of podocytes expressing the target gene, bubble colour indicating expression level, grey line indicating the split between DNA repair and mTORC1 target genes. E: Selected scatterplots of ERCC1 and ERCC4 expression data of single podocytes obtained from living donor (LD), Minimal change disease (MCD), and FSGS biopsies. F: Selected scatterplots of RNA polymerase 1-3 subunit POLR2K and RNA polymerase 2 subunit POLR2J of single podocytes obtained from living donor (LD), Minimal change disease (MCD), and FSGS biopsies. G: Representative immunofluorescence staining of synaptopodin (SNP), γH2A.X, and Draq5 in sections of human MCD and FSGS biopsies, yellow dotted line indicating nuclear border, scalebar indicating 2 µm with quantification of γH2A.X foci per podocyte nuclear area and per podocyte nucleus in human MCD and FSGS biopsies (n=4, 4 glomeruli per sample, 5 podocytes per glomerulus). All violin plots indicate median (black) and upper and lower quartile (gray), *p ≤ 0,05, ***p ≤ 0,001, ****p ≤ 0,0001.

Furthermore, the striking podocyte phenotype upon DNA damage accumulation prompted us to investigate DNA damage repair genes in different glomerular diseases. When analysing data of the human renal cDNA biobank (ERCB) which contains cDNA samples of kidney biopsies, we detected alterations in the expression of DNA repair genes in glomerular lysates of different nephropathies compared to controls (Fig. S7; Tbl. S1), with the strongest effect in focal segmental glomerulosclerosis (FSGS) and the mildest in minimal change disease (MCD). Hence, we investigated the expression of DNA repair genes in podocytes of human FSGS biopsies, minimal change disease (MCD) and healthy controls of a single nucleus RNA sequencing data set (50) (Fig. 5C). Indeed, we detected an upregulation in *Ercc1-8* genes, all involved in damage recognition, DNA unwinding, and damage excision (Fig. 5D, E & S8) in FSGS podocytes. This upregulation indicates that podocytes increase their machinery to repair DNA damage in the presence of FSGS. Vice versa, the expression of several subunits of RNA polymerase 2 and the shared subunit of RNA polymerases 1-3, POL2RK, were virtually lost in human podocytes of both MCD and FSGS samples (Fig 5D, F & S8). These expression data indicate a predominance of damage recognition and strand excision in FSGS biopsies with the potential of downregulation / stalling of RNA synthesis. Targets of the mTORC1 pathway served as an internal control, depicting increased expression with little changes in the percentage of podocytes expressing these genes (Fig. 5D).

To further elucidate this observation of perturbed DNA damage recognition, we investigated the accumulation of DNA damage in podocyte nuclei of human biopsies of FSGS compared to MCD patients using γH2A.X immunofluorescence staining. Indeed, we detected a marked increase in podocyte-specific nuclear γH2A.X foci in FSGS glomeruli (Fig. 5G) corresponding to increased DNA double-strand breaks and thus suggestive of an involvement of DNA damage accumulation in human glomerular disease with marked podocyte damage and loss.

In line with this, using an expression quantitative trait locus (eQTL) analysis, we identified single nucleotide polymorphisms (SNPs) associated with alterations in the expression of DNA repair genes (Table S2 – S4) in sclerotic glomerular diseases that could be targets of interest for future investigations and also for therapeutic modulation.

## Discussion

To which extent glomerular epithelial cells are subject to transcription stress is currently unclear. Elucidating this subject is of particular interest for podocytes, terminally differentiated cells with a limited ability to self-renew (7). These cells have to maintain a functional genome for the organism’s entire life-span, specifically important in the later stages of life. For the first time, our study reveals the occurrence of DNA damage foci in individual podocytes under healthy conditions, indicating a need for constant DNA maintenance and repair and an increase of DNA damage in aged podocytes. Furthermore, we show that this accumulation of DNA damage results in the newly established principle of transcriptional stalling and a shift to the expression of shorter transcripts in glomeruli (35). Recent studies have indicated that the aged podocyte phenotype is in part characterized by senescence. Transcriptomic analysis detected differential expression of senescence-associated genes in aged podocytes (61) coinciding with increased gene silencing. This senescence phenotype was shown to be driven by GSK3 beta overexpression and inhibition via lithium could improve podocyte aging in mice and decrease podocyturia in humans (62). This fits well to the interplay of DNA damage and GSK3 beta activation, as GSK3 beta was shown to be involved DNA damage repair in cancer and that its nuclear translocation was, in part, dependent on p53 activation and nuclear translocation (63–65). A potential causal chain for the senescent phenotype could, therefore, be the activation of GSK3 beta signaling due to DNA damage accumulation in aged podocytes.

The response to transcriptomic stress in podocytes was different from the pattern reported in other cell types, as podocytes activated mTORC1 signaling upon DNA damage. The interplay of DNA damage, its repair, and the mTOR pathway has been a subject of numerous studies, specifically in the field of cancer biology. These studies indicated that mTORC1 signaling is inhibited upon DNA damage in a TSC, Sestrin or AKT-dependent manner (18)(19)(20). Strikingly, we observed that podocytes both *in vitro* and *in vivo* reacted to endogenous accumulation or exogenous infliction of DNA damage with activation of the mTORC1 pathway. Herein lies a fundamental difference to past reports and a potential disease mechanism as numerous studies have depicted the importance of a tight regulation of mTORC1 for podocyte health and the deleterious effects of both overactivation and repression in disease (10, 24, 26, 44, 59, 66). Since mTORC1 activation occurs in *Ercc1* pko podocytes at 9 weeks, a time point of no overt podocyte loss, mild ultrastructural differences, and significant accumulation of DNA damage foci, our data indicate a link between transcription stress and mTORC1 activation in podocytes. This activation occurs via activation of DNA-PK, a nuclear serine/threonine protein kinase.

A growing body of evidence suggests that mTORC1 activity reduces the capacity of successful DNA repair (21, 22), e.g. through ribosomal S6 kinase (S6K)-dependent phosphorylation of E3 ubiquitin-protein ligase RNF168 (23). However, upon podocyte depletion, remaining podocytes on the glomerular tuft counteract the loss of neighbouring cells through mTORC1-mediated hypertrophy (10) which impairs proper DNA repair. This fact is underlined by podocyte-specific *Tsc1* knockout animals displaying mTORC1 hyperactivation. These animals accumulate a significant amount of DNA damage foci already at 4 weeks of age. We, therefore, hypothesize that the interplay of DNA damage and mTORC1 signaling can lead to a downward spiral: accumulation of DNA damage in podocytes triggers mTORC1 activation which further aggravates insufficient DNA maintenance leading to excessive podocyte loss. This cascade could potentially be modulated through well-timed mTORC1 inhibition as suggested by the reduction of glomerulosclerosis in rapamycin-treated *Ercc1* -/delta and *Ercc1* pko animals or by upregulation of DNA repair mechanisms to alleviate accumulated transcription stress.

Only recently the importance of DNA repair has sparked larger interest in the field of podocyte biology. The description of glomerular abnormalities in human syndromes caused by mutations in DNA repair genes is scarce due to their frequently lethal phenotypes (67, 68). First evidence of ERCC1 being an essential protein in kidney health was recently provided through the identification of *ERCC1* variants causing proximal tubular dysfunction as well as glomerular disease underlined by albuminuria in the reported patients (69). Data from whole body *Ercc1* null mice with correction of the liver phenotype also indicate a role for glomerular health with *Ercc1*-deficient animals rapidly dying of renal failure (70). Beyond that, there is evidence for podocyte involvement in human syndromes caused by mutations of DNA repair genes with patients exhibiting proteinuria and nephrotic syndrome (55, 71, 72). Consistently, gene deletion of DNA repair endonuclease co-factor *Ercc1* in podocytes caused proteinuria and FSGS in our mouse model.

The same holds true for factors involved in other forms of DNA maintenance such as the KEOPS complex (14, 73), KAT5 (74), and SMPDL3b in podocytes, playing a role in DNA damage recognition (75). Recently, these processes were shown to extend towards epigentic effects, as HDAC deletion in podocytes resulted in DNA damage accumulation, senescence, cell cycle reentry, and FSGS (76). Itoh and co-workers linked proteinuric kidney disease present in patients with hypertension and diabetes to DNA double-strand breaks and methylation in the promotor region of the slit diaphragm protein nephrin, and established an association to DNA double-strand breaks in glomeruli of patients suffering from IgA nephropathy (15, 16). Our analysis adds considerably to this body of evidence, as we identified multiple factors of DNA maintenance to be transcriptionally altered in glomeruli and single podocytes of different renal diseases involving pronounced podocyte damage and loss, such as MCD and FSGS. Interestingly, we found increased expression of DNA repair proteins in focal segmental glomerulosclerosis (FSGS), suggesting that increased transcription stress may contribute to the loss of podocytes in renal disease. This is in line with altered expression of endonuclease ERCC4 in IgA nephropathy (15). Our study complements these data suggesting that loss of especially transcription-coupled nucleotide excision repair capacity, which removes lesions from the template DNA strands of actively transcribed genes, may contribute to the progression of some types of glomerular disease and aging. The importance of DNA damage repair in podocyte homeostasis is consistent with the role in other postmitotic cell types and particularly apparent in neurodegenerative pathologies typical for DNA repair deficiency syndromes (5). Repairing transcription-blocking lesions might thus play a pivotal role in podocytes as post-mitotic cells that need to maintain the integrity of transcribed genes during the entire lifespan of the organism. Even hepatocytes of *Ercc1* -/delta mice, usually characterized by high self-renewing potential, showed a considerable block of transcription, indicative of transcription-coupled mechanisms being stalled (50, 77). Potential sites in the genome that may be used to alter glomerular damage repair were already identified in our expression quantitative trait locus (eQTL) analysis in FSGS patients. This is of particular interest since there seems to be a broad interplay between gene products exerting functions beyond their canonical pathways in genome maintenance (78). Likewise, decreased expression of DNA repair genes in a subset of patients or the accumulation of DNA damage through the aging process may lead to increased mTORC1 activation in podocytes, thereby rendering a subgroup of patients vulnerable to the development of glomerular disease and scarring.

In conclusion, we identified efficient DNA damage repair as an essential stress response mechanism for podocyte homeostasis and established a link between DNA damage accumulation and mTORC1 signaling. Furthermore, the presented study identifies transcription stress as a hallmark of podocyte aging, loss, and glomerular disease, suitable for precision medicine approaches.

## Author contributions

FB, AMM, LB, RAH, VGP, DM, MNW, GC, MR, DF, GGS, SK, MTL, WB, VN, PM performed experiments; FB, LB, AMM, GC, RAH, VGP, MNW, DF, MTL, WB, MGS, VN analysed data; FB, AMM, VGP, GG, MD, PTB, CDC, MK, MGS, MK, TBH, BSche, TB, BSchu, CEK conceived experiments, FB, VGP, BSchu, CEK wrote the manuscript, all authors revised the manuscript, all authors agreed on the publication in the presented state.

## Funding

Fabian Braun received funding from the Else-Kröner-Fresenius-Stiftung: Else-Kröner-Memorial Stipendium / 2021_EKMS.26

Amrei M. Mandel was partially funded by the Else Kröner-Fresenius-Stiftung and the Eva Luise und Horst Köhler Stiftung – Project No: 2019_KollegSE.04.

Linda Blomberg was supported by the Faculty of Medicine, University of Cologne: Koeln Fortune Program KF 12/2020.

Victor Puelles received support from the Deutsche Forschungsgemeinschaft: Clinical Research Unit / CRC1192, and the Bundesministerium für Bildung und Forschung eMed Consortia / Fibromap) and the Novo Nordisk Foundation (NNF21OC0066381).

Georgia Chatzinikolaou is supported by ELIDEK grant agreement No 1059, funded from the Hellenic Foundation for Research and Innovation (HFRI) and the General Secretariat for Research and Technology (GSRT).

Gisela G. Slaats was supported by an EMBO long-term fellowship ALTF 475-2016 and the Fritz Thyssen Foundation project 10.20.1.012MN.

Martijn Dollé received support the National Institute of Health (NIH)/National Institute of Aging (NIA) (AG17242)

Paul T. Brinkkoetter received support from the Deutsche Forschungsgemeinschaft: Clinical research unit: KFO 329, BR 2955/8-1, and the Bundesministerium für Bildung und Forschung STOP-FSGS 01GM1901E

Matthew G. Samspon is supported by NIH grants RO1 DK108805, DK119380, and RC2 DK122397

George A. Garinis was supported by ELIDEK grant 631, funded from the Hellenic Foundation for Research and Innovation (HFRI).

The George M. O’Brien Michigan Kidney Translational Core Center was funded by NIH/NIDDK grant 2P30-DK-081943.

Bernhard Schermer received support by the Deutsche Forschungsgemeinschaft: Clinical research unit KFO 329 SCHE 1562/7-1.

Thomas Benzing received support from the Deutsche Forschungsgemeinschaft: Clinical research unit: KFO 329, BE 2212/23-1 & 2212/24-1, and the Bundesministerium für Bildung und Forschung STOP-FSGS 01GM1901E

Christine Kurschat received support by the Deutsche Forschungsgemeinschaft: Clinical research unit KFO 329 KU 1539/5-1.

## Supporting information

Supplemental Tabl 1

Supplemental Tabl 2

Supplemental Tabl 3

Supplemental Tabl 4

## Acknowledgments

The excellent technical expertise of Martyna Brütting, Melanie Schaper, Anja Obser, Ilka Edenhofer and Bhawani Nagarajah is gratefully acknowledged. The DNA damage response inhibitors used in this study were a kind gift of Ron Jachimowicz. All schematics in figures were created with BioRender.com. The ERCB-KFB was supported by the Else Kröner-Fresenius Foundation. We also thank all participating centres of the European Renal cDNA Bank -Kröner-Fresenius biopsy bank (ERCB-KFB) and their patients for their cooperation. Active members at the time of the study see (N. Shved et al., Scientific reports 7, 8576 (2017)). The <u>Nephrotic</u> <u>Syndrome Study Network</u> (NEPTUNE) is part of the <u>Rare Diseases Clinical Research</u> <u>Network</u> (RDCRN), which is funded by the National Institutes of Health (NIH) and led by the <u>National Center for Advancing Translational Sciences</u> (NCATS) through its <u>Division of Rare</u> <u>Diseases Research Innovation</u> (DRDRI). NEPTUNE is funded under grant number U54DK083912 as a collaboration between NCATS and the <u>National Institute of Diabetes and Digestive and</u> <u>Kidney Diseases</u> (NIDDK). Additional funding and/or programmatic support is provided by the University of Michigan, NephCure Kidney International and the Halpin Foundation. RDCRN consortia are supported by the RDCRN Data Management and Coordinating Center (DMCC), funded by NCATS and the <u>National Institute of Neurological Disorders and Stroke</u> (NINDS) under U2CTR002818.

Funding support for the DMCC is provided by NCATS and the National Institute of Neurological Disorders and Stroke (NINDS).

Members of the Nephrotic Syndrome Study Network (NEPTUNE) Members of the Nephrotic Syndrome Study Network (NEPTUNE) NEPTUNE Enrolling Centers

*Cleveland Clinic, Cleveland, OH*: K Dell*, J Sedor**, M Schachere^#^, J Negrey^#^

*Children’s Hospital, Los Angeles, CA*: K Lemley*, S Tang^#^

*Children’s Mercy Hospital, Kansas City, MO*: T Srivastava*, S Morrison^#^ *Cohen Children’s Hospital, New Hyde Park, NY:* C Sethna*, M Pfaiff ^#^ *Columbia University, New York, NY:* P Canetta*, A Pradhan^#^

*Emory University, Atlanta, GA:* L Greenbaum*, C Wang**, E Yun^#^

*Harbor-University of California Los Angeles Medical Center:* S Adler*, J LaPage^#^ *John H. Stroger Jr. Hospital of Cook County, Chicago, IL:* A Athavale*, M Itteera *Johns Hopkins Medicine, Baltimore, MD:* M Atkinson*, T Dell^#^

*Mayo Clinic, Rochester, MN:* F Fervenza*, M Hogan**, J Lieske*, V Chernitskiy^#^

*Montefiore Medical Center, Bronx, NY:* F Kaskel*, M Ross*, P Flynn^#^

*NIDDK Intramural, Bethesda MD:* J Kopp*, J Blake^#^

New York *University Medical Center,* New York*, NY:* L Malaga-Dieguez*, O Zhdanova**, F Modersitzki^#^, L Pehrson^#^

Stanford *University,* Stanford*, CA:* R Lafayette*, B Yeung^#^

*Temple University,* Philadelphia*, PA:* I Lee*, S Quinn-Boyle^#^

*University Health Network* Toronto: H Reich *, M Hladunewich**, P Ling^#^, M Romano^#^

*University of Miami, Miami, FL:* A Fornoni*, C Bidot^#^

*University of Michigan, Ann Arbor, MI:* M Kretzler*, D Gipson*, A Williams^#^, C Klida^#^

*University of North Carolina, Chapel Hill, NC:* V Derebail*, K Gibson*, A Froment^#^, S Kelley^#^ *University of Pennsylvania,* Philadelphia*, PA:* L Holzman*, K Meyers**, K Kallem^#^, A Swenson^#^ *University of Texas Southwestern,* Dallas*, TX:* K Sambandam*, K Aleman^#^, M Rogers^#^

*University of Washington,* Seattle*, WA:* A Jefferson*, S Hingorani**, K Tuttle**^§^, M Bray ^#^, E Pao^#^, A Cooper^#§^

*Wake Forest University Baptist Health, Winston-Salem, NC:* JJ Lin*, Stefanie Baker^#^

*Data Analysis and Coordinating Center*: M Kretzler*, L Barisoni**, C Gadegbeku**, B Gillespie**, D Gipson**, L Holzman**, L Mariani**, M Sampson**, J Sedor**, J Zee**, G Alter, H Desmond, S Eddy, D Fermin, M Larkina, S Li, S Li, CC Lienczewski, T Mainieri, R Scherr, A Smith, A Szymanski, A Williams.

*Digital Pathology Committee:* Carmen Avila-Casado (University Health Network, Toronto), Serena Bagnasco (Johns Hopkins University), Joseph Gaut (Washington University in St Louis), Stephen Hewitt (National Cancer Institute), Jeff Hodgin (University of Michigan), Kevin Lemley (Children’s Hospital of Los Angeles), Laura Mariani (University of Michigan), Matthew Palmer (University of Pennsylvania), Avi Rosenberg (Johns Hopkins University), Virginie Royal (University of Montreal), David Thomas (University of Miami), Jarcy Zee (University of Pennsylvania) Co-Chairs: Laura Barisoni (Duke University) and Cynthia Nast (Cedar Sinai).

*Principal Investigator; **Co-investigator; #Study Coordinator

§Providence Medical Research Center, Spokane, WA

## Figures and Figure Legends

## Supplementary Information

**Fig S1:**
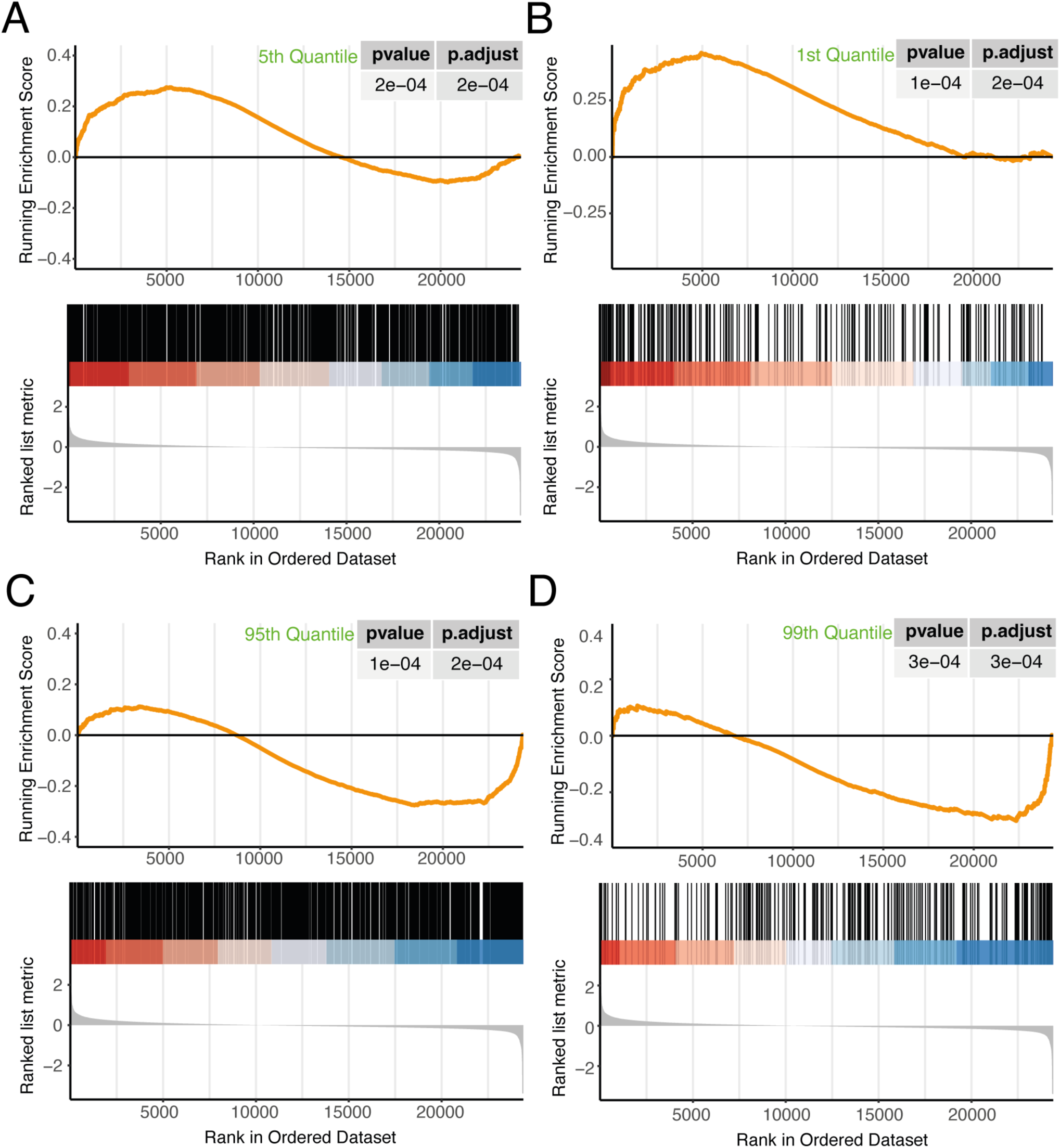
GSEA of gene classes according to transcript length. Shown are the analyses for the shortest 5 % (top left), shortest 1 % (top right), longest 5% (bottom left), and longest 1% (bottom right) of genes. The bottom of each panel shows the log2 fold changes of the microarray data in a ranked order. The top panels show the running enrichment score as an orange line. The smallest 1% (normalized enrichment score (NES) of 2.01) and 5% (NES of 1.37) of genes are significantly more enriched in the upregulated genes, while the longest 1% (NES of -1.69) and 5% (NES of -1.42) of genes are significantly enriched in the downregulated genes in the comparison *Ercc1* -/delta 14 weeks vs. WT 14 weeks.

**Figure S2:**
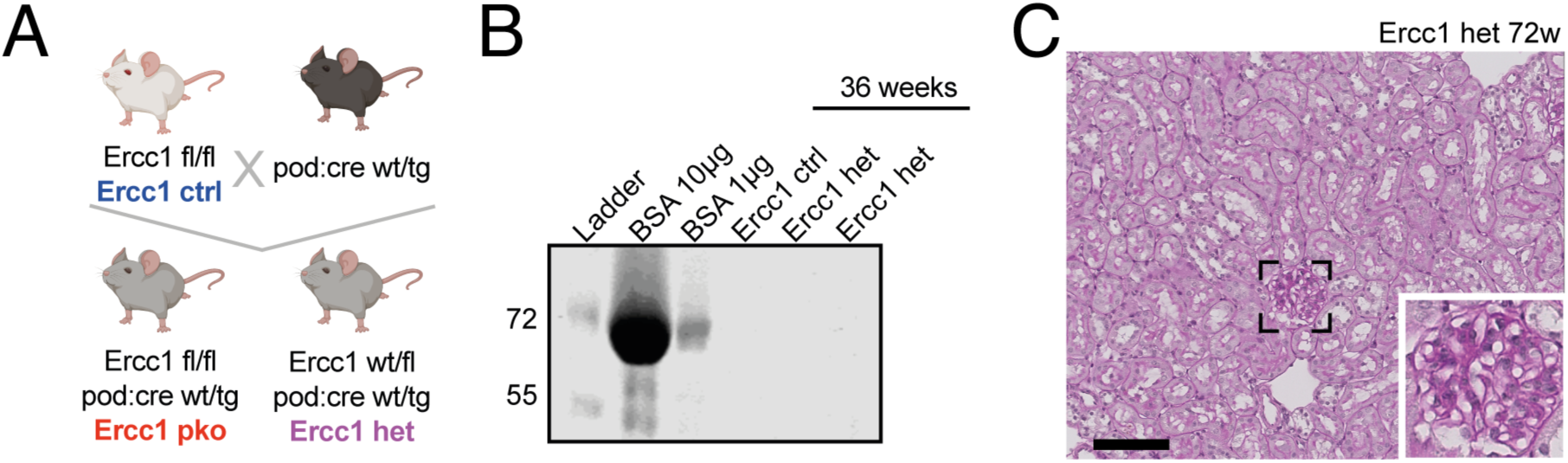
A: Breeding scheme for homozygous and heterozygous podocyte-specific Ercc1 pko mice. B: Representative Coomassie blue staining of *Ercc1* ctrl and wt/pko (het) urine at 36 weeks of age; bovine serum albumin (BSA) was loaded as reference. C: Representative Periodic Acid Schiff (PAS) staining of *Ercc1* wt/pko (het) kidney at 72 weeks of age, scalebar: 100 µm.

**Figure S3:**
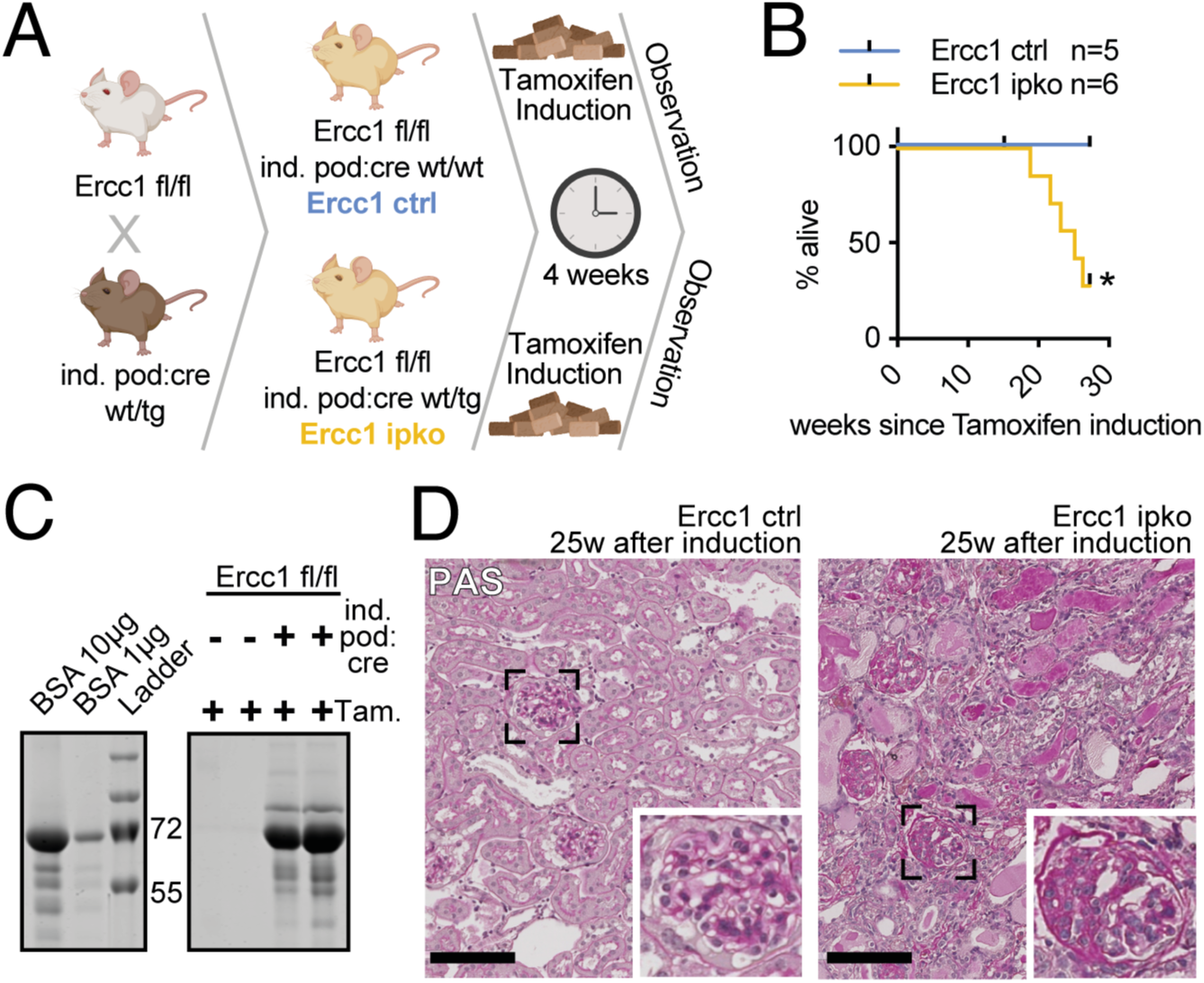
A: Breeding and induction scheme for homozygous inducible podocyte-specific *Ercc1* ko mice (ipko). B: Kaplan-Meyer curve depicting survival of *Ercc1* ctrl and ipko mice (Mantel-Cox test). C: Representative Coomassie blue staining of *Ercc1* ctrl and ipko urine 18 weeks after induction with tamoxifen; bovine serum albumin (BSA) was loaded as reference (n=6). D: Representative Periodic Acid Schiff (PAS) staining of *Ercc1* ctrl and ipko mice 25 weeks after induction with tamoxifen (n=6). *p ≤ 0,05, scalebar: 100µm.

**Figure S4:**
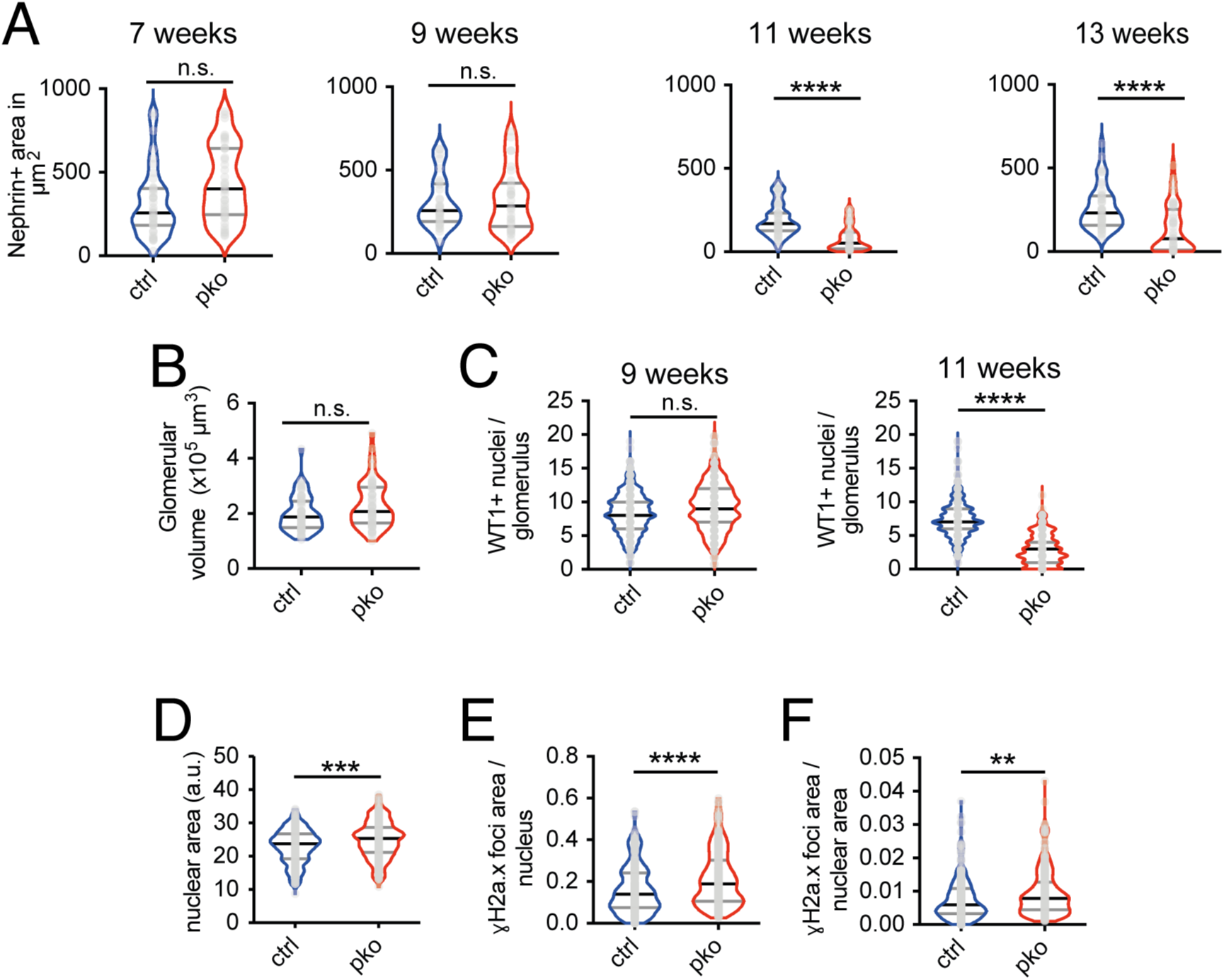
A: Quantification of nephrin positive area in µm^2^ of *Ercc1* ctrl and pko kidneys at 7, 9, 11, and 13 weeks of age, n=4, 10 glomeruli per sample. B: Quantification of glomerular volume of 9-week-old *Ercc1* ctrl and pko kidneys, n = 5, 10 glomeruli per sample. C: Quantification of WT+ nuclei per glomerulus of *Ercc1* ctrl and pko kidneys at 9 and 11 weeks of age, n = 4, ≥50 glomeruli per group. D: Quantification of podocyte nuclear area of 9-week-old *Ercc1* ctrl and pko kidneys, n = 5, 10 glomeruli per sample, 5 podocytes per glomerulus. E: Quantification of γH2A.X foci area per podocyte nucleus of 9-week-old *Ercc1* ctrl and pko kidneys, n = 5, 10 glomeruli per sample, 5 podocytes per glomerulus. F: Quantification of γH2A.X foci area per podocyte nuclear area of 9-week-old *Ercc1* ctrl and pko kidneys, n = 5, 10 glomeruli per sample, 5 podocytes per glomerulus. All violin plots indicate median (black) and upper and lower quartile (gray), **p ≤ 0,01, ***p ≤ 0,001, ****p ≤ 0,0001.

**Figure S5:**
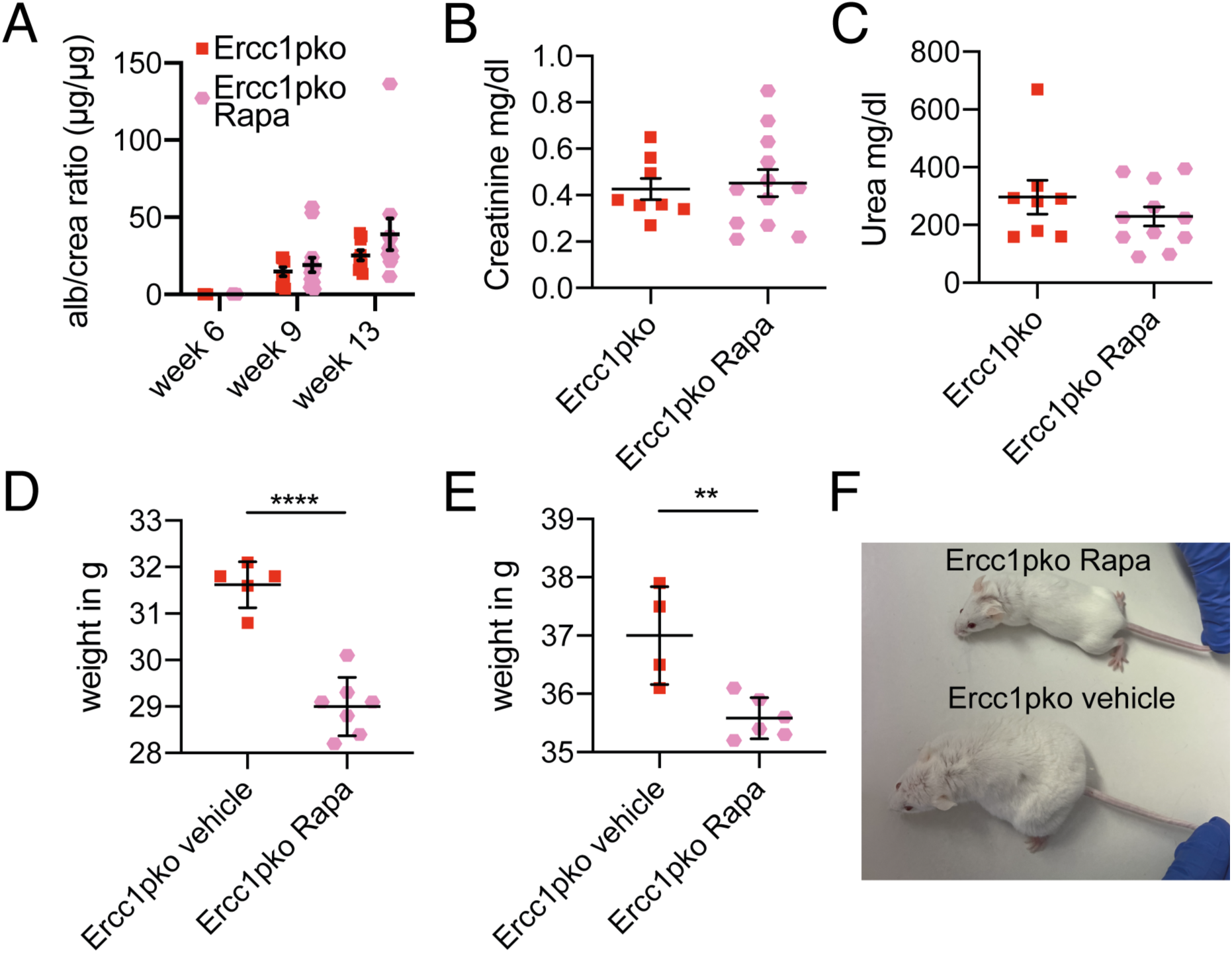
A: urinary albumin/creatinine analysis; B: serum creatinine analysis; C: serum urea analysis and weight analysis (D: female; E: male) of *Ercc1* pko mice treated with vehicle (Ercc1pko) or rapamycin (Ercc1pko Rapa). F: Representative image of rapamycin and vehicle treated *Ercc1* pko mice at 13 weeks of age depicting edema in the vehicle treated animal. All scatterplots indicate mean plus 95% confidence interval, **p ≤ 0,01, ****p ≤ 0,0001, scalebars: 50µm.

**Figure S6:**
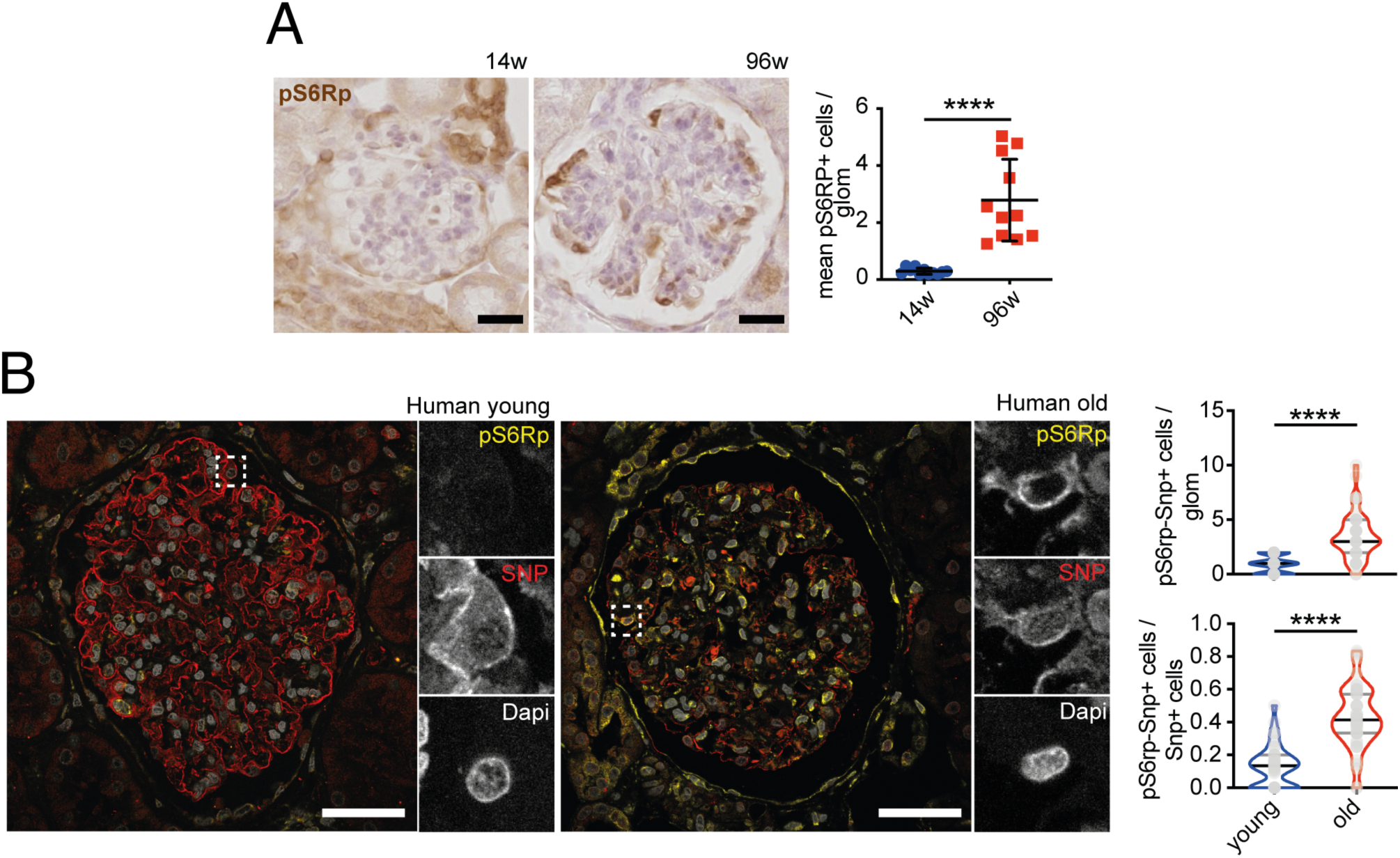
A: Representative immunohistochemistry staining of pS6RP in sections of murine young and aged wildtype kidneys with quantification of pS6RP-positive cells per glomerulus, scalebar indicating 25 µm, n = 11, 50 glomeruli per sample. B: Representative immunofluorescence staining of SNP, pS6RP, and DAPI in sections of young and old human tumor nephrectomy kidneys with quantification of SNP and pS6RP double positive cells per glomerulus and per total SNP positive cells, scalebar indicating 10 µm, n=≥4, 10 glomeruli per sample. Scatterplot depicting mean and 95% confidence interval, all violin plots indicating median (black) and upper and lower quartile (gray), ****p ≤ 0,0001.

**Figure S7:**
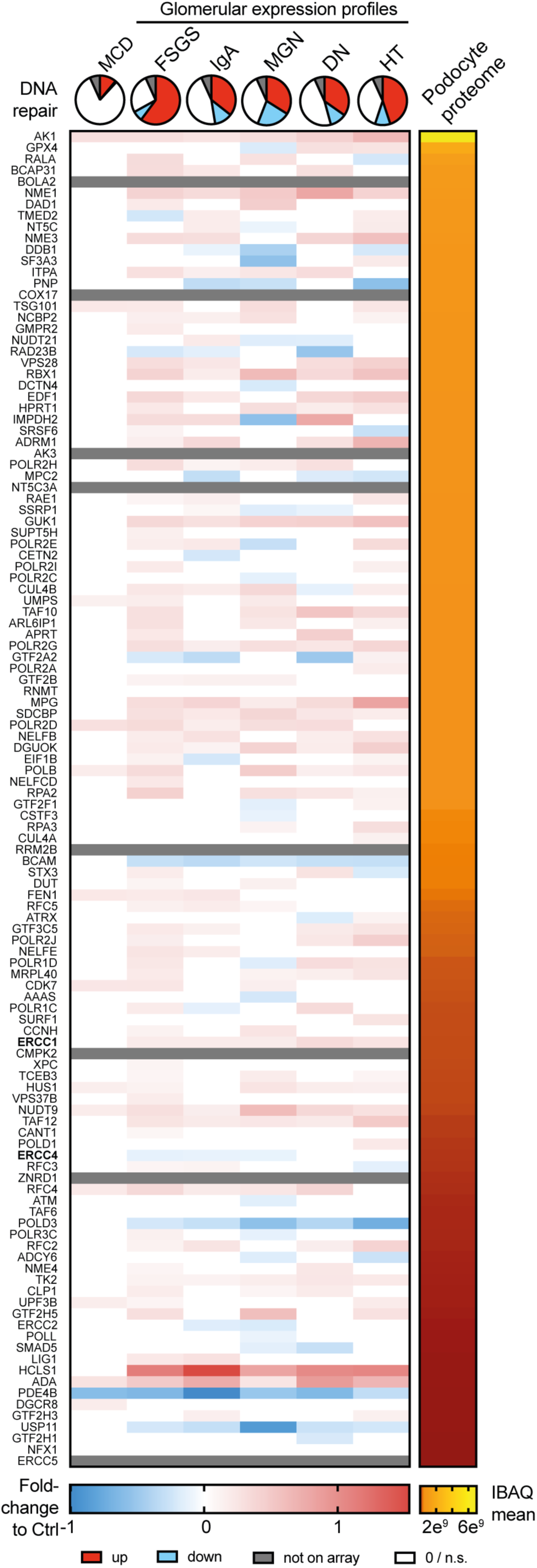
Expression profile of 118 hallmark DNA repair genes in MCD, FSGS, IgA nephropathy (IgA), membranous nephropathy (MGN), diabetic nephropathy (DN), and hypertension (HT) glomeruli compared to controls depicted as parts of whole and single genes in heatmaps. Genes ranked by their protein abundance (Intensity-based absolute quantification - IBAQ) in murine podocyte proteome analysis (41).

**Figure S8:**
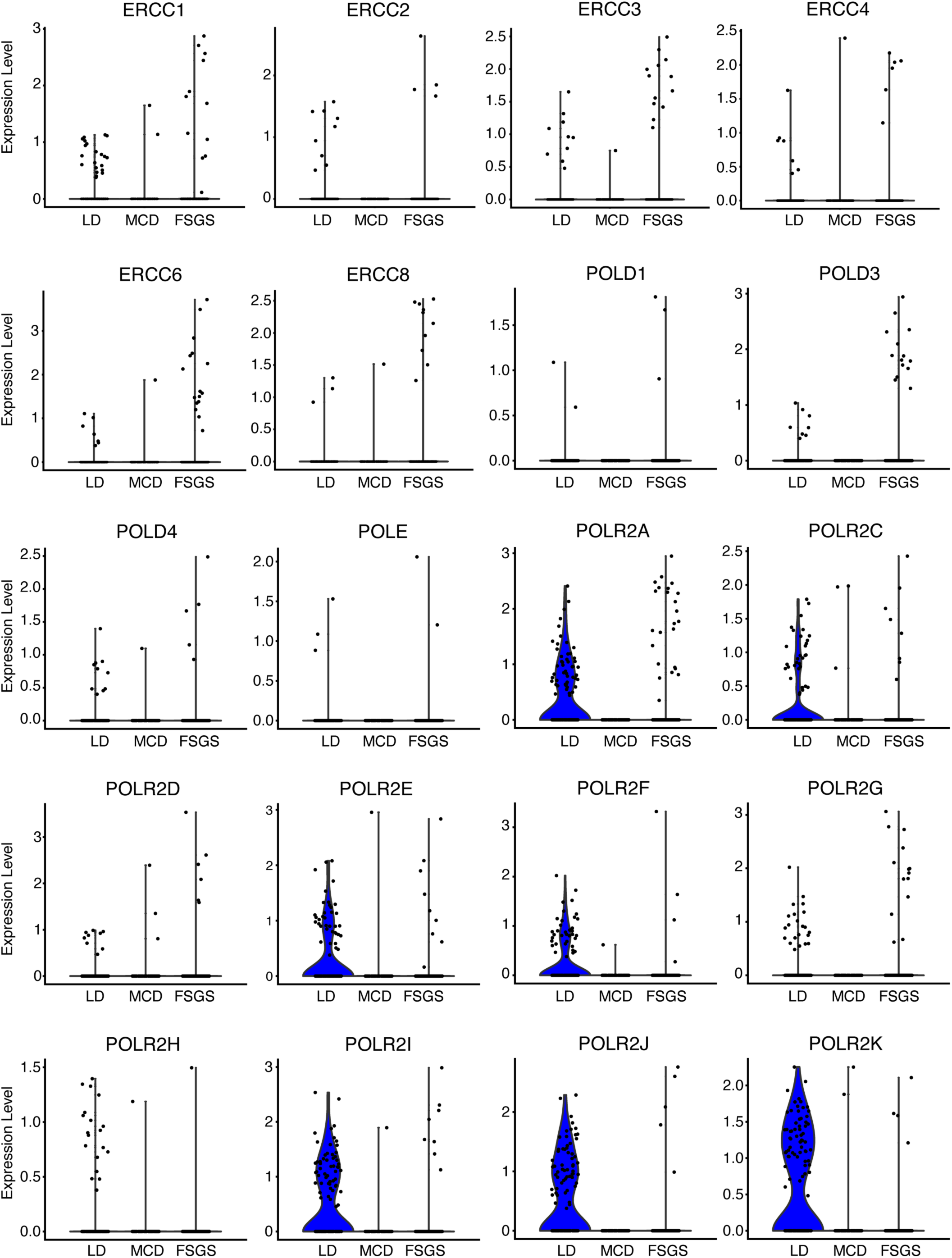
Scatterplots of ERCC family and RNA polymerase subunit expression data of single podocytes obtained from living donor (LD), Minimal change disease (MCD), and FSGS biopsies.

**Table S1: Gene expression analysis of ERCB for Hallmark DNA Repair and Nucleotide Excision Repair Genes**

**Table S2: Clinical characteristics of FSGS patients**

**Table S3: eQTL analysis of DNA repair genes in FSGS patients** Ensg: ensemble gene ID; FDR: false discovery rate, PIP: posterior inclusion probability, AF: allele frequency, beta: expression difference to reference allele

**Table S4: Full eQTL analysis of DNA repair genes in FSGS patients**

